# Dysregulated balance of D- and L-amino acids modulating glutamatergic neurotransmission in severe spinal muscular atrophy

**DOI:** 10.1101/2024.10.22.619645

**Authors:** Amber Hassan, Raffaella di Vito, Tommaso Nuzzo, Matteo Vidali, Maria Jose Carlini, Shubhi Yadav, Hua Yang, Adele D’Amico, Xhesika Kolici, Valeria Valsecchi, Chiara Panicucci, Giuseppe Pignataro, Claudio Bruno, Enrico Bertini, Francesco Errico, Livio Pellizzoni, Alessandro Usiello

**Author notes:** Alessandro Usiello, Ph.D.: Department of Environmental, Biological and Pharmaceutical Sciences and Technologies, University of Campania “Luigi Vanvitelli”, Via A. Vivaldi, 43, 81100 Caserta, Italy, and CEINGE Biotecnologie Avanzate, Naples, Italy. These authors contributed equally to this work (Amber Hassan, Raffaella di Vito, Tommaso Nuzzo).

## Abstract

Spinal muscular atrophy (SMA) is a neuromuscular disorder caused by reduced expression of the survival motor neuron (SMN) protein. In addition to motor neuron survival, SMN deficiency affects the integrity and function of afferent synapses that provide glutamatergic excitatory drive essential for motor neuron firing and muscle contraction. However, it is unknown whether deficits in the metabolism of excitatory amino acids and their precursors contribute to neuronal dysfunction in SMA. To address this issue, we measured the levels of the main neuroactive D- and L-amino acids acting on glutamatergic receptors in the central nervous system of SMNΔ7 mice as well as the cerebrospinal fluid (CSF) of SMA patients of varying severity before and after treatment with the SMN-inducing drug Nusinersen. Our findings reveal that SMN deficiency disrupts glutamate and serine metabolism in the CSF of severe SMA patients, including decreased concentration of L-glutamate, which is partially corrected by Nusinersen therapy. Moreover, we identify dysregulated L-glutamine to L-glutamate conversion as a shared neurochemical signature of altered glutamatergic synapse metabolism that implicates astrocyte dysfunction in both severe SMA patients and mouse models. Lastly, consistent with a correlation of higher CSF levels of D-serine with better motor function in severe SMA patients, we show that daily supplementation with the NMDA receptor co-agonist D-serine improves neurological deficits in SMNΔ7 mice. Altogether, these findings provide direct evidence for dysregulation of D- and L-amino acid metabolism linked to glutamatergic neurotransmission in severe SMA and have potential implications for treating this neurological disorder.

## Introduction

Spinal muscular atrophy (SMA) is the most common genetic cause of infant mortality and is characterized by the progressive degeneration of spinal motor neurons and skeletal muscle atrophy (1–3), with increasing evidence of multiorgan pathology (3, 4). SMA is caused by homozygous mutations in the *Survival Motor Neuron 1* (*SMN1)* gene (5), while the paralogue gene *SMN2* - which is present in variable copy numbers - can only partially compensate for *SMN1* loss due to a single nucleotide change affecting its pre-mRNA splicing (6). SMA patients are classified into three main clinical groups (Types 1, 2 and 3) based on the age of onset and severity of the disease, which inversely correlate with the number of *SMN2* copies and the SMN protein levels (1).

Three different therapeutic approaches aimed at increasing SMN expression through splicing modulation or viral-mediated gene replacement have been approved for the treatment of SMA (4). Among these, Nusinersen (Sрinrаzа) - the first FDA-approved drug for the treatment of SMА (7) - is an antisense oligonucleotide administered intrathecally that corrects the splicing of *SMN2* pre-mRNA, increasing the concentration of functional SMN protein (4, 6, 8). Clinical data show that Nusinersen induces remarkable improvement of motor function in SMA patients, especially when treated pre-symptomatically (9). Despite significant advances in SMA therapy, however, the current consensus is that neither Nusinersen nor other treatments can cure the disease (4, 8, 10). In particular, there are still unmet needs to address the incomplete correction of disease symptoms and the variability in clinical response to treatment. One of the major limitations is the absence of validated targets whose modulation could increase the clinical benefit of SMN-inducing drugs through combinatorial therapy (11, 12). In this regard, the identification of neurometabolic markers linked to disease severity and correlated with functional outcomes after treatment would be crucial for explaining differences in clinical responses and guiding discovery of new therapies, including nutritional support. However, our current understanding of the biochemical and neurochemical abnormalities associated with SMA pathology and their specific response to SMN-inducing therapies is very limited.

Studies in model organisms revealed that cell-autonomous deficits in motor neurons alone cannot account for the disease phenotype and implicated dysfunction of excitatory neuronal networks that control motor neuron output as important players of SMA pathophysiology (13–17). The best characterized example is the dysfunction and loss of glutamatergic synapses from proprioceptive sensory neurons to motor neurons, which have emerged as some of the earliest manifestations of the disease in SMA mice (13, 14, 17, 18). The resulting reduction in afferent glutamatergic neurotransmission causes downregulation of Kv2.1 channels and decreased firing of SMA motor neurons, contributing to impaired muscle contraction and motor dysfunction (17). Consistent with SMA motor neurons suffering from reduced excitatory drive, increasing neuronal activity and glutamate signaling on motor neurons improve motor function in animal models of SMA (13, 15, 17, 19). Importantly, recent studies have expanded the detrimental effects of SMN deficiency in neuronal networks beyond hypoglutamatergic signaling to include noradrenergic and serotonergic neurotransmission (20, 21). However, it is unknown whether deficits in the metabolism of excitatory amino acids contribute to synaptic dysfunction in SMA sensory-motor circuits.

L-glutamate (L-Glu) is the most important excitatory amino acid in the central nervous system (CNS) (22). It plays a pivotal role in orchestrating neurodevelopmental processes, synaptic transmission, and plasticity within the brain and spinal cord through stimulation of ionotropic (NMDA, AMPA, and kainate) and metabotropic (mGlu) receptors (23–28). Besides neurotransmission, L-Glu controls fundamental cellular pathways, including non-essential amino acid synthesis and energy metabolism by directly regulating α-ketoglutarate levels and, in turn, the Krebs cycle (29). The closely related dicarboxylic amino acid L-aspartate (L-Asp) and its D-enantiomer derivative D-Asp are also known to act as primary agonists of NMDARs (26, 30–32) and mGluR5 (33), while D-serine (D-Ser) functions in glutamatergic signaling by acting as an NMDAR co-agonist (34, 35). Lastly, L-glutamine (L-Gln) and L-Ser play a primary role in modulating the synthesis of L-Glu and D-Ser and, together with L- Asp, are involved in essential cellular processes including energy homeostasis, ammonium recycling, redox balance and in the biosynthesis of amino acids, nucleotides, and membrane lipids (36).

Despite their critical roles for neuronal function, possible alterations in the physiological levels of neuroactive amino acids in the CNS of either SMA patients or animal models have not been investigated. Here, we addressed this question by performing a comprehensive neurotransmitter profiling in the spinal cord and brain of severe SMA mice as well as in the cerebrospinal fluid (CSF) of SMA patients across the spectrum of disease severity. We also investigated the impact of Nusinersen therapy on the CSF levels of excitatory amino acids and their precursors implicated in glutamatergic neurotransmission. Overall, our findings provide direct evidence for dysregulation of amino acid metabolism linked to glutamatergic neurotransmission that may contribute to motor dysfunction in severe SMA.

## Results

### SMN deficiency perturbs glutamate and serine metabolism in the CSF of severe SMA patients

We performed a real-world, retrospective study to determine the effects of SMN deficiency on the concentration of neuroactive amino acids and their precursors in the CSF of untreated SMA patients across the disease-severity spectrum using HPLC (Fig.1A). This analysis included CSF from SMA1 (*n* = 34), SMA2 (*n* = 22), and SMA3 (*n* = 17) patients as well as age-matched control subjects (*n* = 7) whose clinical and demographic features are presented in Table 1.

**Figure 1.**
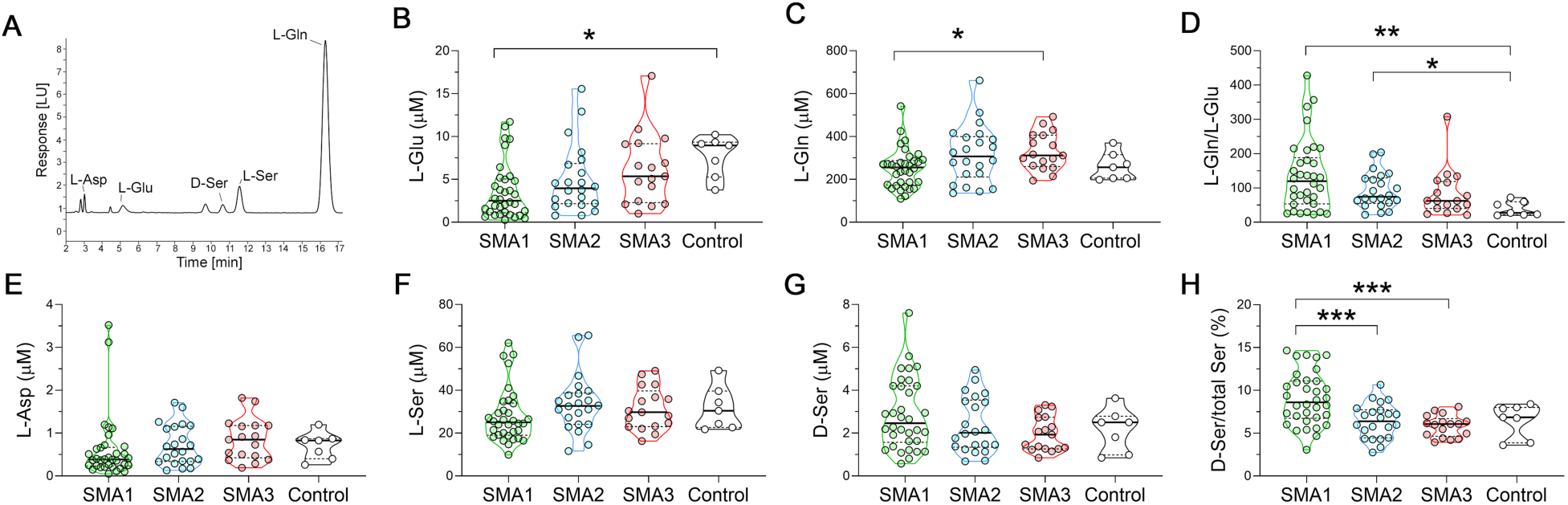
Levels of neuroactive amino acids in the CSF of SMA patients and control individuals. **(A)** Representative chromatogram showing the peaks of L-aspartate (L-Asp), L-glutamate (L-Glu), D-serine (D-Ser), L-serine (L-Ser), and L-glutamine (L-Gln) in CSF of SMA1 patients. **(B-H)** Levels of L-Glu **(B)**, L-Gln **(C)**, L-Gln/L-Glu ratio **(D)**, L-Asp **(E)**, L-Ser **(F)**, D-Ser **(G)** and D-Ser/total Ser percentage ratio **(H)** in the indicated cohorts of SMA1, SMA2 and SMA3 patients as well as control individuals. Data are shown as violin plots representing the median with interquartile range (IQR). \**P* < 0.05, \*\**P* < 0.01(Mann-Whitney test with Bonferroni’s correction). Dots represent values from each individual analyzed.

**Table 1.** Demographic and clinical characteristics of naïve SMA subjects and controls enrolled in the study.

| Demographic and clinical information | Controls (n=7) | SMA1 (n=34) | SMA2 (n=22) | SMA3 (n=17) |
| --- | --- | --- | --- | --- |
| Sex (M:F, %) | 43 : 57 | 40 : 60 | 41: 59 | 37 : 63 |
| Age, years | 7.0 (4.0-12.0) | 3.0 (0.7-4.9) | 5.5 (2.0-11.8) | 13.7 (7.6-17.2) |
| SMN copy number (2:3:4, %) |  | 91:9:0 | 10:90:0 | 21.4: 64.3 14.3 |
| BMI |  | 13.4 (12.4-15.3) | 17.9 (15.9-19.8) | 19.4 (16.2-22.1) |
| CHOP-INTEND |  | 14 (3-21) | - | - |
| HFMSE |  | - | 9.5 (6.5-15) | 52 (35-57) |
| Gastrostomy (Yes:No, %) |  | 51 : 49 | 0 :100 | 0 :100 |
| NIV (Yes:No, %) |  | 41 : 59 | 33 : 67 | 8 : 92 |
| Tracheostomy (Yes:No, %) |  | 34 :66 | 0 : 100 | 0 :100 |

Data are expressed as percentage frequencies or median (IQR). Abbreviations: BMI = Body Mass Index; CHOP INTEND = Children's Hospital Of Philadelphia Infant Test Of Neuromuscular Disorders; HFMSE = Hammersmith Functional Motor Scale Expanded; NIV= Non-invasive ventilation.

Using the non-parametric Kruskal-Wallis test, we found significant L-Glu and L-Gln changes in the CSF of SMA patients (L-Glu, *P* = 0.007; L-Gln, *P* = 0.023) (Fig. 1B,C; Suppl. Table 1). Importantly, further *post-hoc* comparisons highlighted a significant reduction of L-Glu in the CSF of SMA1 patients compared to controls (Fig. 1B; Suppl. Table 1). The L-Gln to L-Glu ratio was also significantly affected in SMA patients (*P* = 0.005; Kruskal-Wallis) (Fig. 1D; Suppl. Table 1). Accordingly, we found increased L-Gln/L-Glu ratio in both SMA1 and SMA2 patients compared to controls (respectively, *P* = 0.006 and *P* = 0.036; Mann-Whitney with Bonferroni correction) (Fig. 1D; Suppl. Table 1). In addition, the D-Ser to total Ser ratio – which is a reliable index of D-Ser metabolism (37) – showed a significant difference among clinical conditions (*P* < 0.001; Kruskal-Wallis) (Fig. 1H and Suppl. Table 1). The following *post-hoc* analysis revealed an increased D-Ser/total Ser ratio in SMA1 patients compared to those with milder forms of SMA (*P* < 0.001; Fig 1H; Suppl. Table 1). The Kruskal-Wallis test showed significant changes in L-Asp levels in SMA patients (*P* = 0.037) (Fig. 1E; Suppl. Table 1), and the CSF levels of D-Asp were below the detection limit of our HPLC settings (0.01 pmol). Lastly, ANCOVA analysis performed on natural log-transformed data to evaluate the potential confounding effects of age and sex indicated that only variations in the levels of L-Glu (*P* = 0.006) and the L-Gln/L-Glu ratio (*P* = 0.003) were significantly associated with the clinical condition. These findings show that SMN deficiency leads to significantly decreased concentrations of L-Glu in the CSF of severe SMA1 patients and to altered L-Gln to L-Glu conversion in the CSF of both SMA1 and SMA2 patients.

We next investigated whether the CSF concentrations of amino acids within each SMA patient type were associated with age or motor function assessed by CHOP-INTEND (SMA1) or HFMSE (SMA2 and SMA3) clinical assays. Non-parametric Spearman’s correlation analysis revealed that the levels of L-Glu and L-Gln as well as the L-Gln/L-Glu ratio showed no significant correlation with age or clinical parameters in SMA1 patients (Table 2). In contrast, D-Ser levels and the D-Ser/total Ser ratio were negatively correlated with age (Table 2; Suppl Fig. 1) and positively correlated with CHOP-INTEND (Table 2; Suppl. Fig. 2). Since we also found a negative correlation between age and CHOP-INTEND (Table 2; Suppl. Fig. 3), we performed multivariate linear regression analysis considering age as a confounding factor. This analysis did not confirm the significance of the association between D-Ser or D-Ser/total Ser ratio with CHOP-INTEND (respectively, *P* = 0.074 and *P* = 0.203), highlighting the putative influence of age on this correlation. In SMA2 patients, statistical analysis showed a significant negative correlation between D-Ser levels and the D-Ser/total Ser ratio with age (Table 2; Suppl. Fig. 1). There were also positive correlations of D-Ser levels and the D-Ser/total Ser ratio with the HFMSE score (Table 2; Suppl. Fig. 2), which failed to reach statistical significance probably due to the small sample size. Similar to SMA1 patients, age and motor function were negatively correlated in SMA2 individuals (Table 2; Suppl Fig. 3). In SMA3 patients, we found a negative correlation of the D-Ser/total Ser ratio with age but not with the HFMSE score (Table 2; Suppl Fig. 1 and 2). There was also no correlation between age and the HFMSE score (Table 2; Suppl. Fig. 3). Interestingly, no significant associations occurred between either D-Ser levels or the D-Ser/total Ser ratio and age in control subjects (Table 2; Suppl Fig. 1), consistent with the possibility that age-dependent decrease in D-Ser levels is specific to the disease state. However, the limited sample size of controls might affect this result. The confounding effects of age notwithstanding, these results highlight a potential correlation between greater D-Ser levels and the D-Ser/total Ser ratio and better motor function, which is especially apparent in severe SMA patients.

**Figure 2.**
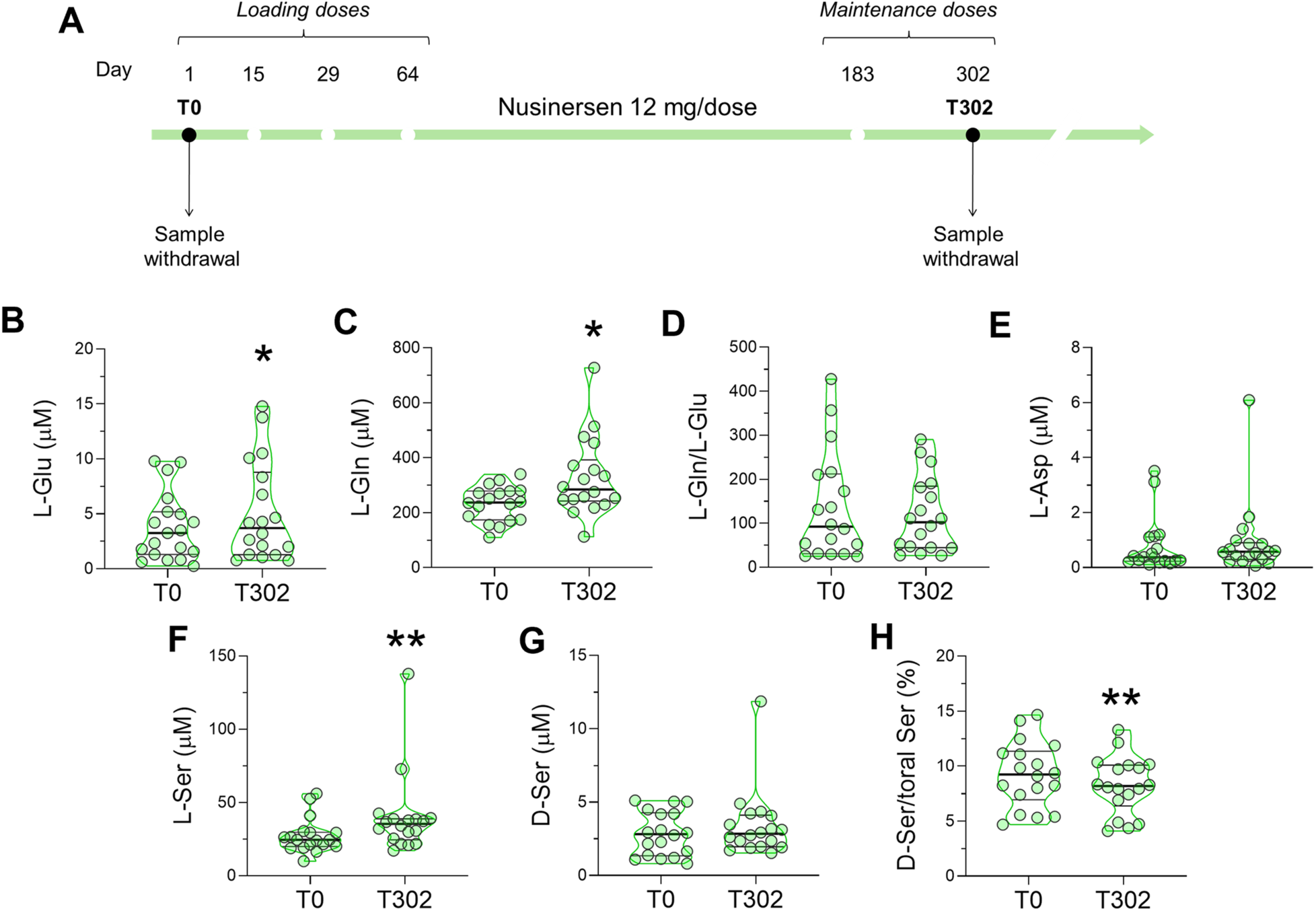
Effect of Nusinersen on the levels of neuroactive amino acids in the CSF of SMA1 patients. **(A)** Schematic representation of the timeline of intrathecal Nusinersen administration and CSF collection in SMA patients. **(B-H)** Levels of L-glutamate (L-Glu) **(B)**, L-glutamine (L-Gln) **(C)**, L-glutamine/L-glutamate (L-Gln/L-Glu) ratio **(D)**, L-aspartate (L-Asp) **(E)**, L-serine (L-Ser) **(F)**, D-serine (D-Ser) **(G)** and D-serine/total serine (D-Ser/total Ser) percentage ratio **(H)** in the CSF of SMA1 patients before treatment (T0, n=18) and at the time of the sixth (T302, n=18) injection of Nusinersen. \**P* < 0.05, \*\**P* < 0.01, compared to T0 (Wilcoxon matched-pairs signed ranks test). Data are shown as violin plots representing the median with interquartile range (IQR). Dots represent values from individual SMA1 patients.

**Table 2.** Correlation between amino acids and clinical or demographic variables.

|  |  | CHOP-INTEND/<br>HFMSE | L-Gln | L-Glu | L-Gln/<br>L-Glu | L-Asp | L-Ser | D-Ser | D-Ser/<br>total Ser |
| --- | --- | --- | --- | --- | --- | --- | --- | --- | --- |
| SMA1 | Age | $\rho=-0.540$<br>**$P=0.001$**<br>n=32 | $\rho=-0.026$<br>$P=0.888$<br>n=32 | $\rho=-0.086$<br>$P=0.641$<br>n=32 | $\rho=0.171$<br>$P=0.349$<br>n=32 | $\rho=0.067$<br>$P=0.714$<br>n=32 | $\rho=-0.044$<br>$P=0.811$<br>n=32 | $\rho=-0.428$<br>**$P=0.015$**<br>n=32 | $\rho=-0.720$<br>**$P<0.001$**<br>n=32 |
| | CHOP-INTEND | - | $\rho=0.07$<br>$P=0.971$<br>n=32 | $\rho=0.196$<br>$P=0.281$<br>n=32 | $\rho=-0.243$<br>$P=0.180$<br>n=32 | $\rho=0.233$<br>$P=0.199$<br>n=32 | $\rho=0.116$<br>$P=0.528$<br>n=32 | $\rho=0.377$<br>**$P=0.034$**<br>n=32 | $\rho=0.452$<br>**$P=0.009$**<br>n=32 |
| SMA2 | Age | $\rho=-0.638$<br>**$P=0.002$**<br>n=20 | $\rho=0.131$<br>$P=0.571$<br>n=21 | $\rho=-0.034$<br>$P=0.882$<br>n=21 | $\rho=0.076$<br>$P=0.743$<br>n=21 | $\rho=0.073$<br>$P=0.752$<br>n=21 | $\rho=-0.199$<br>$P=0.388$<br>n=21 | $\rho=-0.470$<br>**$P=0.032$**<br>n=21 | $\rho=-0.577$<br>**$P=0.006$**<br>n=21 |
| | HFMSE | - | $\rho=0.049$<br>$P=0.837$<br>n=20 | $\rho=-0.032$<br>$P=0.892$<br>n=20 | $\rho=0.077$<br>$P=0.747$<br>n=20 | $\rho=-0.328$<br>$P=0.158$<br>n=20 | $\rho=0.165$<br>$P=0.488$<br>n=20 | $\rho=0.378$<br>$P=0.100$<br>n=20 | $\rho=0.381$<br>$P=0.097$<br>n=20 |
| SMA3 | Age | $\rho=0.338$<br>**$P=0.238$**<br>n=14 | $\rho=0.178$<br>$P=0.543$<br>n=14 | $\rho=-0.231$<br>$P=0.427$<br>n=14 | $\rho=0.270$<br>$P=0.350$<br>n=14 | $\rho=-0.292$<br>$P=0.311$<br>n=14 | $\rho=-0.257$<br>$P=0.375$<br>n=14 | $\rho=-0.464$<br>$P=0.095$<br>n=14 | $\rho=-0.692$<br>**$P=0.006$**<br>n=14 |
| | HFMSE | - | $\rho=0.201$<br>$P=0.491$<br>n=14 | $\rho=0.020$<br>$P=0.946$<br>n=14 | $\rho=0.029$<br>$P=0.922$<br>n=14 | $\rho=0.174$<br>$P=0.551$<br>n=14 | $\rho=0.024$<br>$P=0.934$<br>n=14 | $\rho=0.082$<br>$P=0.781$<br>n=14 | $\rho=0.117$<br>$P=0.690$<br>n=14 |
| Controls | Age | - | $\rho=0.286$<br>$P=0.535$<br>n=7 | $\rho=-0.321$<br>$P=0.482$<br>n=7 | $\rho=0.429$<br>$P=0.337$<br>n=7 | $\rho=0.071$<br>$P=0.879$<br>n=7 | $\rho=-0.750$<br>$P=0.052$<br>n=7 | $\rho=-0.714$<br>$P=0.071$<br>n=7 | $\rho=-0.484$<br>$P=0.294$<br>n=7 |

Non-parametric Spearman's rho coefficients ( $\rho$ ) and related $P$ -values ( $P$ ) are indicated. Significant $P$ -values are shown in bold. Abbreviations: CHOP-INTEND = Children's Hospital of Philadelphia Infant Test of Neuromuscular Disorders; HFMSE = Hammersmith Functional Motor Scale Expanded.

### Nusinersen modulates glutamate, glutamine, and serine levels in the CSF of severe SMA patients

We next sought to determine the effects of Nusinersen treatment on the CSF levels of neuroactive amino acids and whether it could normalize the alterations observed in untreated, early-onset SMA patients. For this longitudinal analysis, we used a subgroup of SMA patients (*n* = 18 SMA1, *n* = 17 SMA2, and *n* = 14 SMA3) for whom CSF samples were available both prior to (T0) and 302 days after (T302) initiation of treatment, corresponding to the maintenance phase of Nusinersen therapy (Figure 2A; Table 3).

**Table 3.** Clinical and demographic characteristics of SMA patients treated with Nusinersen.

| Demographic and clinical information | SMA1 (n=18) | SMA2 (n=17) | SMA3 (n=14) |
| --- | --- | --- | --- |
| Sex (M:F, %) | 39 : 61 | 30 : 70 | 30 : 70 |
| Age | 2.7 (0.7-4.9) | 5.5 (3.1-11.8) | 13 (9-16) |
| SMN copy number (2:3:4, %) | 94:6:0 | 94:6:0 | 23:63:14 |
| BMI | 13.2 (11.7-14.6) | 18.2 (16.3-19.8) | 19.4 (16.2-22.1) |
| CHOP-INTEND | 6.5 (3-17) |  |  |
| HFMSE |  | 9 (8-13) | 55 (36-56) |
| Gastrostomy (Yes:No, %) | 66:34 | 0:100 | 0/100 |
| NIV (Yes:No, %) | 39: 61 | 35:65 | 8:92 |
| Tracheostomy (Yes:No, %) | 33: 67 | 0:100 | 0:100 |

Data are expressed as percentage frequencies or median (IQR). Abbreviations: BMI = Body Max Index; CHOP INTEND = Children's Hospital Of Philadelphia Infant Test Of Neuromuscular Disorders; HFMSE = Hammersmith Functional Motor Scale Expanded; NIV= Non-invasive ventilation.

The non-parametric Wilcoxon matched-pairs test revealed that Nusinersen therapy significantly increased the levels of L-Glu (*P* = 0.035) and L-Gln (*P* = 0.025) in the CSF of SMA1 patients relative to the corresponding drug-free baseline (Fig. 2B, C; Suppl. Table 2). Moreover, Nusinersen administration significantly increased L-Ser levels (*P* = 0.004) and decreased the D-Ser/total Ser ratio in the CSF of SMA1 patients (*P* = 0.007) (Fig. 2F,H; Suppl. Table 2). In SMA2 patients, we did not observe significant changes in the concentration of neuroactive amino acids after Nusinersen treatment (Suppl. Fig. 4; Suppl. Table 2). In SMA3 patients, Nusinersen treatment decreased the CSF levels of several amino acids, which differs from its effects in both SMA1 and SMA2 patients. Accordingly, we found lower levels of L-Glu (*P* = 0.006) resulting in a small increase in the L-Gln/L-Glu ratio (*P* = 0.019) as L-Gln levels were comparable between groups (*P* = 0.064) (Suppl. Fig. 5A-C; Suppl. Table 2). There were also reduced levels of L-Ser (*P* = 0.019), D-Ser (*P* = 0.016), and L-Asp (*P* = 0.038) in Nusinersen-treated SMA3 patients relative to baseline (Suppl. Fig. 5D-F; Suppl. Table 2). Overall, these results reveal that Nusinersen-dependent upregulation of SMN modulates the metabolism of L-Glu, L-Gln, and L-Ser while decreasing the D-Ser/total Ser ratio in the CSF of SMA1 patients, partially counteracting the dysregulation of amino acids associated with the severe form of the disease prior to treatment.

We next investigated the association of changes in amino acid levels with demographic and clinical parameters of SMA patients following 302 days of Nusinersen treatment. Non-parametric Spearman’s correlation analysis highlighted a positive correlation between the D-Ser/total Ser ratio and CHOP-INTEND but the lack of significant correlation with age in Nusinersen-treated SMA1 patients (Suppl. Fig. 6; Table 4). However, when controlling for age by multivariate linear regression, the association between D-Ser/total Ser and CHOP-INTEND was lost (*P* = 0.136). As expected for the early-onset severe form of the disease, we found a negative correlation between age and CHOP-INTEND in SMA1 patients (Table 4). In Nusinersen-treated SMA2 patients, we found a positive correlation of L-Ser and D-Ser levels with HFMSE (Suppl. Figure 7-8; Table 4). Specifically, L-Ser and D-Ser levels were negatively correlated with age, which was also negatively correlated with HFMSE (Table 4). However, only age but not L-Ser or D-Ser remained significantly associated with HFMSE after multivariate linear regression analysis (L-Ser: *P* = 0.308; D-Ser: *P* = 0.809).

**Table 4.** Correlation between amino acids and clinical or demographic variables in SMA1, SMA2 and SMA3 patients before and after treatment with Nusinersen.

| Patients | Therapy phase | Parameter | CHOP-INTEND / HFMSE | L-Gln | L-Glu | Gln/Glu | L-Asp | L-Ser | D-Ser | D-Ser/tot Ser |
| --- | --- | --- | --- | --- | --- | --- | --- | --- | --- | --- |
| SMA1<br>(N=18) | T0 | Age | $\rho=-0.636$<br>**$P=0.005$** | $\rho=0.084$<br>$P=0.741$ | $\rho=0.071$<br>$P=0.779$ | $\rho=0.018$<br>$P=0.945$ | $\rho=0.237$<br>$P=0.343$ | $\rho=-0.005$<br>$P=0.984$ | $\rho=-0.344$<br>$P=0.162$ | $\rho=-0.649$<br>**$P=0.004$** |
| | | CHOP-INTEND | - | $\rho=0.113$<br>$P=0.656$ | $\rho=0.292$<br>$P=0.240$ | $\rho=-0.326$<br>$P=0.187$ | $\rho=0.325$<br>$P=0.188$ | $\rho=0.257$<br>$P=0.304$ | $\rho=0.408$<br>$P=0.093$ | $\rho=0.389$<br>$P=0.110$ |
| | T302 | Age | $\rho=-0.735$<br>**$P=0.001$** | $\rho=0.141$<br>$P=0.576$ | $\rho=0.189$<br>$P=0.453$ | $\rho=0.001$<br>$P=0.997$ | $\rho=0.352$<br>$P=0.152$ | $\rho=-0.005$<br>$P=0.984$ | $\rho=-0.257$<br>$P=0.303$ | $\rho=-0.364$<br>$P=0.137$ |
| | | CHOP-INTEND | - | $\rho=-0.303$<br>$P=0.237$ | $\rho=-0.225$<br>$P=0.386$ | $\rho=-0.039$<br>$P=0.881$ | $\rho=-0.184$<br>$P=0.479$ | $\rho=-0.139$<br>$P=0.596$ | $\rho=0.264$<br>$P=0.306$ | $\rho=0.528$<br>**$P=0.030$** |
| SMA2<br>(N=17) | T0 | Age | $\rho=-0.511$<br>**$P=0.036$** | $\rho=0.020$<br>$P=0.940$ | $\rho=0.025$<br>$P=0.926$ | $\rho=-0.065$<br>$P=0.804$ | $\rho=0.040$<br>$P=0.877$ | $\rho=-0.207$<br>$P=0.425$ | $\rho=-0.415$<br>$P=0.098$ | $\rho=-0.471$<br>$P=0.056$ |
| | | HFMSE | - | $\rho=0.102$<br>$P=0.696$ | $\rho=-0.119$<br>$P=0.648$ | $\rho=0.217$<br>$P=0.404$ | $\rho=-0.437$<br>$P=0.079$ | $\rho=0.053$<br>$P=0.840$ | $\rho=0.233$<br>$P=0.369$ | $\rho=0.177$<br>$P=0.496$ |
| | T302 | Age | $\rho=-0.845$<br>**$P<0.001$** | $\rho=-0.422$<br>$P=0.091$ | $\rho=-0.329$<br>$P=0.197$ | $\rho=-0.145$<br>$P=0.579$ | $\rho=-0.188$<br>$P=0.471$ | $\rho=-0.677$<br>**$P=0.003$** | $\rho=-0.729$<br>**$P=0.001$** | $\rho=-0.218$<br>$P=0.400$ |
| | | HFMSE | - | $\rho=0.402$<br>$P=0.109$ | $\rho=0.330$<br>$P=0.196$ | $\rho=0.028$<br>$P=0.914$ | $\rho=0.098$<br>$P=0.707$ | $\rho=0.522$<br>**$P=0.032$** | $\rho=0.568$<br>**$P=0.017$** | $\rho=0.038$<br>$P=0.884$ |
| SMA3<br>(N=13) | T0 | Age | $\rho=0.337$<br>$P=0.259$ | $\rho=0.206$<br>$P=0.499$ | $\rho=-0.124$<br>$P=0.687$ | $\rho=0.157$<br>$P=0.609$ | $\rho=-0.217$<br>$P=0.476$ | $\rho=-0.248$<br>$P=0.415$ | $\rho=-0.437$<br>$P=0.135$ | $\rho=-0.710$<br>**$P=0.007$** |
| | | HFMSE | - | $\rho=0.210$<br>$P=0.491$ | $\rho=0.017$<br>$P=0.957$ | $\rho=0.039$<br>$P=0.900$ | $\rho=0.210$<br>$P=0.491$ | $\rho=0.033$<br>$P=0.914$ | $\rho=0.110$<br>$P=0.719$ | $\rho=0.110$<br>$P=0.719$ |
| | T302 | Age | $\rho=0.262$<br>$P=0.388$ | $\rho=0.393$<br>$P=0.184$ | $\rho=0.025$<br>$P=0.936$ | $\rho=0.165$<br>$P=0.590$ | $\rho=0.201$<br>$P=0.511$ | $\rho=0.044$<br>$P=0.886$ | $\rho=-0.105$<br>$P=0.734$ | $\rho=-0.746$<br>**$P=0.003$** |
| | | HFMSE | - | $\rho=0.536$<br>$P=0.059$ | $\rho=0.377$<br>$P=0.204$ | $\rho=-0.094$<br>$P=0.761$ | $\rho=0.198$<br>$P=0.517$ | $\rho=0.272$<br>$P=0.368$ | $\rho=0.358$<br>$P=0.230$ | $\rho=0.039$<br>$P=0.901$ |

Non-parametric Spearman's rho coefficients ( $\rho$ ) and related $P$ -values ( $P$ ) are indicated. Significant $P$ -values are shown in bold. Abbreviations: CHOP-INTEND = Children's Hospital of Philadelphia Infant Test of Neuromuscular Disorders; HFMSE = Hammersmith Functional Motor Scale Expanded.

Lastly, statistical analysis showed only a negative correlation of the D-Ser/total Ser ratio with age in Nusinersen-treated SMA3 patients (Table 4). These results show a positive correlation between D-Ser/total Ser and CHOP-INTEND in SMA1, and between L-Ser or D-Ser and HFMSE in SMA2. However, the influence of age on motor function in early-onset SMA patients complicates the interpretation of the observed amino acid variations in relationship to motor improvement.

### Dysregulation of glutamate-glutamine metabolism in the brain and spinal cord of SMA mice

To expand on our investigation of the effects of SMN deficiency on the levels of neuroactive amino acids, we conducted HPLC analysis (Fig.3B) in brain and spinal cord tissues isolated from SMA mice at early (postnatal day 3, P3) and late (P11) symptomatic stages of the disease. SMN depletion did not change the levels of any amino acid tested in the brain or spinal cord from SMNΔ7 mice compared with WT at P3 (Fig. 3C,D; Suppl. Fig. 8; Suppl. Table 3). In contrast, we found a significant increase in the concentration of L-Gln and L-Gln/L-Glu ratio in both the brain and spinal cord of SMNΔ7 mice compared to WT at P11 (Fig 3E,F; Suppl. Fig. 9; Suppl. Table 3). Furthermore, SMNΔ7 mice displayed an increased D-Asp/total Asp ratio in the spinal cord at P11 (Suppl. Fig. 9; Suppl. Table 3).

**Figure 3.**
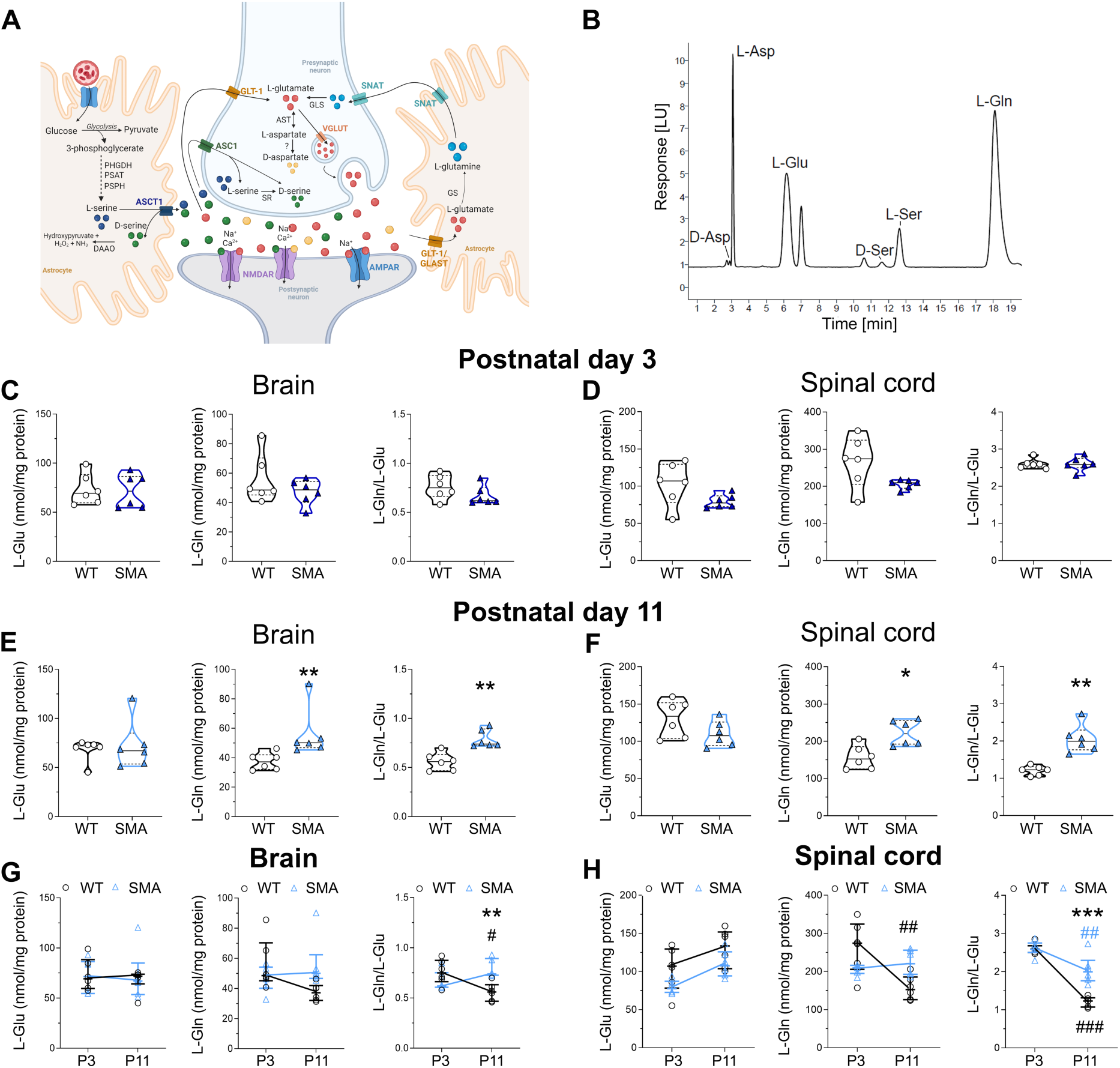
Analysis of neuroactive amino acid levels in the brain and spinal cord of SMNΔ7 mice at early and late symptomatic stage of the disease. **A)** Schematic model of the tripartite glutamatergic synapse showing the main localization of the amino acids analyzed in this study. Image created with BioRender.com (www.biorender.com). Abbreviations: PHGDH: phosphoglycerate dehydrogenase; PSAT: phosphoserine aminotransferase; PSPH: phosphoserine phosphatase; DAAO: D-amino acid oxidase; SR: serine racemase; GLS: glutaminase; GS: glutamine synthetase; ASCT1: alanine, serine, cysteine transporter 1; ASC1: alanine, serine, cysteine transporter 1; GLT-1: glutamate transporter 1; SNAT: sodium-coupled neutral amino acid transporter; GLAST: glutamate aspartate transporter; NMDAR: N-methyl-D-aspartate receptor; AMPAR: alpha-amino-3-hydroxy-5-methyl-4-isoxazolepropionic acid receptor; VGLUT: vesicular glutamate transporter. **B)** Representative chromatogram showing the peaks of L-aspartate (L-Asp), L-glutamate (L-Glu), D-serine (D-Ser), L-serine (L-Ser), and L-glutamine (L-Gln) in the brain homogenate of SMNΔ7 mice. **(C-F)** Levels of L-Glu, L-Gln and L-Gln/L-Glu ratio in the brain and spinal cord of wild type (WT) and SMNΔ7 mice at postnatal day 3 (P3), and P11. The average amounts of amino acids detected were normalized for mg of total proteins. Dots represent values from individual mice. Amino acid levels are expressed as violin plots representing median with interquartile range (IQR) and analyzed by Mann-Whitney test (\**P* < 0.05, \*\**P* < 0.01, compared to age-matched WT mice). (**G,H**) Amino acid levels were also analyzed as two-way ANOVA, followed by Tukey’s multiple comparisons test (\*\**P* < 0.01, \*\**P* < 0.0001, compared to age-matched WT mice; ^##^*P* < 0.01, ^###^*P* < 0.0001, compared to genotype-matched P3 mice). Amino acid levels were shown as scatter dot plots representing median with interquartile range (IQR) while dots represent values from individual mice.

We then used two-way ANOVA followed by Tukey’s *post-hoc* comparisons to further analyze the neurochemical variations during postnatal CNS development in WT and SMNΔ7 mice. This highlighted a physiological, age-dependent drop of the L-Gln/L-Glu ratio in the brain of WT mice that does not occur in SMNΔ7 mice, resulting in a higher L-Gln/L-Glu ratio in SMNΔ7 relative to WT mice at P11 (Fig. 3G; Suppl. Table 4). Similarly, we found that the L-Gln/L-Glu ratio in the spinal cord of SMNΔ7 mice significantly increased relative to WT littermates at P11 (Fig. 3H; Suppl. Table 4). Furthermore, despite significant age-dependent changes between WT and SMNΔ7 mice were found for the D-Ser/total Ser ratio in the brain and the D-Asp/total Asp ratio in the spinal cord, no differences were highlighted between genotypes at each single time point using the Tukey’s multiple comparisons *post-hoc* analysis (Suppl. Fig. 10, Suppl. Table 4).

These findings indicate that SMN deficiency significantly impacts the metabolism of neuroactive amino acids involved in glutamatergic neurotransmission in the brain and spinal cord of SMA mice at the disease end stage. Importantly, the increase in the L-Gln/L-Glu ratio emerges as a conserved signature of neurochemical dysregulation in the CSF of severe SMA patients and the CNS of mouse models.

### D-serine supplementation moderately improves motor function in SMA mice

The results of our neurochemical profiling highlighted the dysregulation of amino acid metabolism acting on glutamatergic neurotransmission as well as a potential correlation between higher D-Ser/total Ser ratio in the CSF and better motor function in severe SMA patients. Since D-Ser is a major co-agonist of NMDARs (34, 35), we sought to investigate the phenotypic effects of increasing D-Ser levels in the CNS of SMA mice.

First, to validate that intraperitoneal (IP) administration of D-Ser increases its levels in the mouse spinal cord, we performed a single injection of vehicle or D-Ser at a dosage of 500 mg/kg in WT mice at P3 followed by HPLC analysis 1 h post-injection (Fig. 4A). As expected (38), we found that the concentration of D-Ser (median [IQR] of nmol/mg of protein, D-Ser = 28.97 [25.29-58.61] vs vehicle *=* 2.81 [2.71-3.09], *P* = 0.0286, Mann-Whitney test) and the D-Ser/total Ser ratio (D-Ser = 51.90 [47.54-67.22] vs vehicle *=* 11.02 [10.34-11.68], *P* = 0.0286) were strongly increased in the spinal cord of D-Ser-injected mice relative to control mice injected with vehicle (Fig. 4A-C).

**Figure 4.**
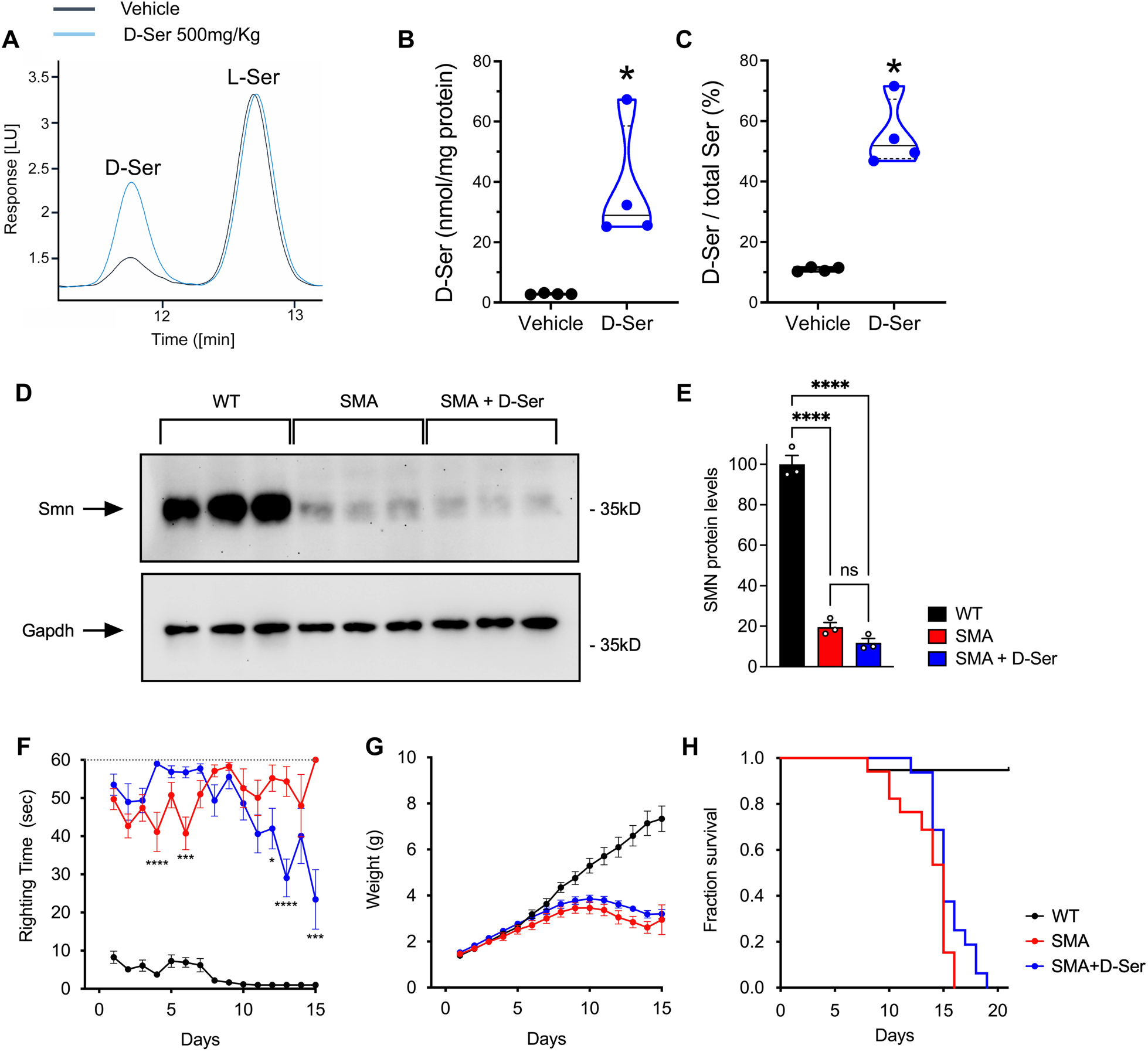
SMN-independent amelioration of motor function by D-serine supplementation in SMA mice. (**A**) Representative chromatogram showing the peaks of D-serine (D-Ser) and L-serine (L-Ser) in the spinal cord homogenate of SMNΔ7 mice after a single injection of vehicle (black line) or 500 mg/kg D-Ser treatment (blu line) at P3. (**B-C**) D-Ser levels and D-Ser/toral Ser ratio in the spinal cord of WT mice treated with a single injection of 500 mg/kg D-serine compared to vehicle-treated mice at P3. \**P* < 0.05, compared to vehicle (Mann-Whitney test) (**D**) Western blot analysis of SMN protein levels in spinal cords from WT and either untreated or D-Ser-treated SMNΔ7 mice at P11. GAPDH was probed as a loading control. Three independent mice per group were analyzed. (**E**) Quantification of SMN levels normalized to GAPDH from the data in (D). Ordinary one-way ANOVA with Šídák’s multiple comparisons test. \*\*\*\**P* < 0.0001, ns: not significant. Values are means and SEM. (**F-H**) Analysis of righting time (F), body weight (G), and survival (H) in WT mice (n=19) and either untreated (n=17) or D-Ser-treated (n=16) SMNΔ7 mice. Two-way ANOVA with Tukey’s multiple comparisons test. \*\*\*\**P* < 0.0001, \*\*\**P* < 0.001, \**P* < 0.05. Values are means and SEM.

Next, we analyzed the effects of daily supplementation of D-Ser (500mg/kg) starting at birth on SMN expression and the phenotype of SMA mice, which included daily measures of weight gain, motor function, and survival. Western blot analysis of spinal cord tissue isolated at P11 from WT and SMA mice either with or without amino acid supplementation demonstrated that D-Ser does not increase SMN protein levels (Fig. 4D,E). Interestingly, phenotypic analysis showed a moderate improvement of motor function assessed by the righting reflex in D-Ser-treated relative to untreated SMA mice (Fig. 4F). This motor benefit appeared in the second postnatal week and was not associated with an increase in weight gain, which was similar for D-Ser-treated and untreated SMA mice (Fig. 4G). Lastly, despite fewer early deaths and some slightly longer-lived mice being noted, treatment with D-Ser did not increase the median survival of SMA mice (Fig. 4H). Overall, these results are consistent with the possibility that increased CNS levels of D-Ser may partially improve motor function in a mouse model of SMA by acting on glutamatergic neurotransmission.

## Discussion

Selective degeneration of motor neurons and skeletal muscle atrophy are hallmarks of SMA in both patients and mouse models (1, 6). However, cell-autonomous deficits in motor neurons alone cannot account for the SMA phenotype and dysfunction of neuronal networks that control motor output has been implicated in disease etiology (6). Accordingly, several studies have shown that reduced excitatory drive to motor neurons (13–17) and metabolomic dysfunction in fundamental intracellular pathways (39) may play a role in SMA pathophysiology. Nevertheless, while a variety of synaptic deficits in motor neuron afferents have been documented (13, 14, 17, 21, 40), whether SMN deficiency directly alters the content of neurotransmitters in the CNS of animal models and SMA patients is unknown. Here, we addressed this issue by investigating changes in the levels of the most abundant excitatory D- and L-amino acids that drive glutamate receptor signaling or act as their immediate precursors in the CNS of SMNΔ7 mice and in the CSF of SMA patients of differing severity before and after Nusinersen therapy.

Our findings reveal that SMN deficiency strongly affects the levels of D- and L-amino acids related to glutamatergic neurotransmission in the CSF of severe SMA patients as well as in the brain and spinal cord of SMNΔ7 mice. Accordingly, we found lower levels of L-Glu in SMA1 patients and a trend toward reduction in SMA2 and SMA3 patients compared to controls. L-Glu reduction also results in a significant increase of the L-Gln/L-Glu ratio in both SMA1 and SMA2 patients compared to control subjects. Interestingly, a prominent increase in the L-Gln/L-Glu ratio also occurs in the brain and spinal cord of SMNΔ7 mice at late symptomatic stages. Thus, deregulated L-Gln to L-Glu conversion emerges as a neurochemical signature of altered glutamatergic metabolism shared by severe SMA patients and animal models that may contribute to impaired excitatory neurotransmission in the disease state.

The SMA-related changes in the L-Gln/L-Glu ratio are suggestive of abnormalities in the glutamate-glutamine cycle, an imbalance of which has been associated with various neurological and psychiatric disorders (41). This cycle involves the functional interaction between neurons and astrocytes in the tripartite glutamatergic synapse where pre- and post-synaptic nerve terminals and perisynaptic astrocytic processes recycle L-Glu for neurotransmission (Fig. 3A). Following synaptic release, L-Glu binds to ionotropic and metabotropic glutamate receptors located post-synaptically and is then taken up by astrocytes via excitatory amino acid transporters to prevent excitotoxicity caused by excessive activation of glutamate receptors (42). In astrocytes, the enzyme glutamine synthetase converts the main part of L-Glu into L-Gln, which is shuttled to the neuron via sodium-coupled neutral amino acid transporters. L-Gln is then deaminated to L-Glu by glutaminase in the mitochondria of neurons and either transferred by vesicular glutamate transporters into synaptic vesicles for the further rounds of neurotransmission or converted in α-ketoglutarate for Krebs cycle activity. Thus, in addition to perturbing glutamatergic neurotransmission, SMN deficiency may also trigger an abnormally increased conversion of L-Glu in α-ketoglutarate into the Krebs cycle as a compensatory mechanism to counteract the pervasive energy failure due to mitochondrial abnormalities associated with SMA (39, 43). Importantly, previous *in vitro* and *in vivo* studies have highlighted several deficits induced by SMN deficiency in SMA astrocytes that may contribute to the dysfunction of the tripartite synapse and the neurochemical alterations observed here (44–47).

Notably, our neurochemical profiling revealed an upregulation of the D-Ser/total Ser ratio in the CSF of SMA1 patients compared to individuals with SMA2 and SMA3, reflecting opposite trends toward decrease or increase of L-Ser and D-Ser levels, respectively. Based on the pharmacological properties of D-Ser as a potent endogenous co-agonist of NMDARs (34, 35), these findings further extend the link between SMN deficiency and dysregulation of amino acid metabolism acting on glutamatergic neurotransmission in severe SMA. As is the case for the glutamate-glutamine cycle described above, an increase in the D-Ser/total Ser ratio points to dysfunction of astrocyte metabolism in SMA1 patients. Accordingly, *de novo* synthesis of L-Ser in the CNS occurs exclusively in astrocytes through the phosphorylated pathway, which employs 3-phosphoglycerate generated by glycolysis; L-Ser is then shuttled to neurons for the biosynthesis of D-Ser catalyzed by serine racemase (SR) (48, 49) (Fig. 3A). Interestingly, our clinical data highlight a positive correlation between D-Ser levels or the D-Ser/total Ser ratio and motor function assessed by CHOP-INTEND in SMA1 patients and a similar trend in SMA2 patients assessed by HFSME. However, the correlation between D-Ser metabolism and motor function should be interpreted cautiously due to the potentially confounding effect of age. Accordingly, age was negatively correlated with D-Ser levels, the D-Ser/total Ser ratio, and motor function in SMA1 and SMA2 patients; and age-adjusted partial correlation and multiple regression analyses did not confirm the association between D-Ser metabolism and clinical scores. The inverse correlation of D-Ser levels or the D-Ser/total Ser ratio with age observed in the CSF of SMA patients and controls is in agreement with previous observation in a large cohort of pediatric individuals (50).

Our longitudinal analysis of the neurochemical composition of the CSF in SMA patients before and after treatment with Nusinersen further supports the role of SMN in regulating the concentration of D- and L-amino acids that modulate glutamatergic neurotransmission, especially in patients affected by the most severe form of the disease. Accordingly, we found that Nusinersen induces a significant increase in the levels of L-Glu and L-Gln as well as upregulation of L-Ser with consequent reduction of the D-Ser/total Ser ratio in the CSF of treated SMA1 patients relative to their drug-free baseline. These findings are consistent with previous untargeted metabolomic studies highlighting the effect of Nusinersen on glutamate metabolism in SMA1 patients (39), which was accompanied by modulation of energy-related Krebs cycle and glutathione metabolism that depend on direct L-Glu supply. Although the mechanisms driving the observed neurochemical variations remain unclear, these results link Nusinersen-dependent SMN upregulation to an improved astrocyte-dependent metabolism of amino acids related to glutamatergic neurotransmission and energy metabolism, which are strongly affected in severe SMA (51, 52). Interestingly, and in contrast to SMA1 patients, Nusinersen did not affect the levels of neuroactive amino acids in the CSF of SMA2 patients and slightly decreased the concentration of L-Glu, L-Asp, L-Ser, D-Ser as well as the L-Gln/L-Glu ratio in SMA3 patients. The opposite effects of Nusinersen on the CSF concentrations of L-Glu and L-Ser in SMA1 and SMA3 patients confirm that the impact of SMN upregulation on amino acid metabolism follows distinct biochemical trajectories depending on SMA severity. Lastly, we found a positive correlation of motor function with the D-Ser/total Ser ratio in SMA1 patients and with the levels of L-Ser or D-Ser in SMA2 patients after 302 days of Nusinersen treatment. As is the case for the analysis in naïve SMA patients, however, it remains unclear whether these neurochemical differences are directly associated with motor improvement because age is a significant confounding factor in these correlations. Nevertheless, it is important to highlight that L-Ser supplementation is currently in use for the therapy of the Neu-Laxova syndrome, characterized by severe peripheral malformations and microcephaly (53) and in Phase I clinical trials for the treatment of an inherited form of peripheral neuropathy (54) and ALS (55). Future longitudinal studies in larger cohorts of SMA patients and age-matched healthy individuals will help address the issue.

Given the difficulty in conclusively establishing cause-effect relationships between changes in the CSF levels of neuroactive amino acids and clinical outcomes in SMA patients, we sought to initially address this issue in a preclinical *in vivo* setting by studying the phenotypic effects of D-Ser metabolism modulation in a mouse model of SMA. The focus on D-Ser was prompted by our observations that the D-Ser/total Ser ratio in the CSF of SMA1 patients i) is higher relative to that of SMA2 and SMA3 patients, ii) positively correlates with motor function, and iii) decreases after Nusinersen therapy. If changes in the D-Ser/total Ser ratio are biologically relevant to SMA pathology, two main scenarios can be envisioned that take into consideration the role of D-Ser as a potent agonist of NMDAR (34, 35). On one hand, increased levels of D-Ser may be beneficial and reflect a homeostatic attempt to counteract deficits in glutamatergic neurotransmission at NMDARs through enhanced L-Ser to D-Ser conversion, which is tuned down after Nusinersen treatment enhances L-Glu levels. On the other hand, increased levels of endogenous D-Ser could be harmful to neurons via excitotoxicity – a possibility consistent with previous studies of the motor neuron disease amyotrophic lateral sclerosis (ALS) (56–58). Our results show that systemic administration of D-Ser by IP injection strongly increases D-Ser levels and the D-Ser/total Ser ratio in the mouse CNS as expected from previous studies (38, 59). Interestingly, daily supplementation of D-Ser in severe SMA mice ameliorates motor function while having no effects on weight gain and survival. Furthermore, the motor improvement is independent from SMN whose low expression levels in the spinal cord are unaffected by D-Ser. These findings suggest the possibility that elevated D-Ser levels do not have deleterious but rather beneficial effects on the severe motor phenotype of SMA mice by enhancing glutamatergic neurotransmission at the GluN1 subunit of NMDARs. This interpretation is in agreement with previous studies showing that physical and pharmacological approaches aimed at increasing neuronal activity and NMDAR signaling improve motor function in animal models of SMA (15, 17, 19, 60–62).

In conclusion, our study shows that SMN deficiency disrupts the physiological balance of neuroactive D- and L-amino acids linked to glutamatergic receptors signaling in the CNS of SMA mice and the CSF of severe SMA patients. The resulting defects may compound the deleterious effects associated with the loss of excitatory synapses on motor neurons in spinal sensory-motor circuits as well as interfere more broadly with glutamatergic neuronal networks in the brain of severe SMA patients, including cognitive deficits. Moreover, our findings identify modulation of glutamate and serine metabolism as downstream targets of Nusinersen treatment in SMA patients and support further investigation of pharmacological approaches, such as D-Ser supplementation, aimed at improving glutamatergic neurotransmission deficits for use in combination therapies with SMN-inducing drugs.

## Materials and Methods

### Patients’ characteristics

This is a two-center study (Bambino Gesù Hospital, Rome, Italy; Giannina Gaslini Institute, Genoa, Italy) conducted on seventy-three patients affected by SMA1 (*n* = 34), SMA2 (*n* = 22) and SMA3 (*n* = 17) who received intrathecal treatment with Nusinersen (12 mg) (Table 1). Additionally, seven non-neurological pediatric control subjects aged 2.5-14 years were included in the study (Table 1). The study was approved by the local Ethics Committees of the two Hospitals (2395_OPBG_2021). All participants and/or their legal guardians signed a written informed consent. CSF samples were collected at day 0 (T0; baseline) and day 302 (T302; after 5 Nusinersen injections) and used for detection of amino acids. For SMA1 patients, we collected *n* = 34 CSF samples at T0, and *n* = 18 at T302. For SMA2 patients, we collected *n* = 22 CSF samples at T0, and *n* = 17 at T302. For SMA3 patients, we collected CSF samples from 17 patients at T0, and 14 CSF samples at T302 (Table 3). For longitudinal analysis, we considered the subgroup of SMA patients (*n* = 18 SMA1, *n* = 17 SMA2, and *n* = 14 SMA3) for whom CSF samples were available both prior to (T0) and 302 days after (T302) initiation of treatment, corresponding to the maintenance phase of Nusinersen therapy (Figure 2A; Table 3). All patients were clinically diagnosed and genetically confirmed, and the *SMN2* copy number was also determined. All SMA1 patients, irrespective of age and disease severity, were part of the Expanded Access Programme (EAP) for compassionate use to patients with the infantile form only, which occurred in Italy between November 2016 and November 2017. The overall clinical response of these patients to Nusinersen treatment has previously been reported as part of the full Italian cohort and showed that therapeutic efficacy is related to age and clinical severity at baseline (63, 64). The SMA2 and SMA3 patients have also been reported previously (65).

### Clinical evaluation

Assessment of patients was performed at T0 and T302. At each visit, extensive clinical examination was performed by experienced child neurologists or pediatricians with expertise in SMA, and anthropometric measurements and vital parameters were collected. Patients’ feeding status (oral nutrition, nasogastric tube (NG) or percutaneous gastrostomy), nutritional status postulated by Body Mass Index (BMI), and respiratory function (spontaneous breathing, non-invasive ventilation (NIV) or tracheostomy) were recorded.

For SMA1 patients, five children were younger than 5 months, while all the others were older than 5 months at the beginning of treatment with ages ranging from 6 months to 10 years. Eleven patients had tracheostomy and thirteen were under NIV for <16h/day. Nineteen patients had gastrostomy, and the BMI fell into the underweight range (< 18.5) in all patients. The age of the SMA2 patients included in this study ranged from 9 months to 13.6 years at baseline. Seven of these patients were under NIV, and fourteen patients were in spontaneous breathing. None had tracheostomy or gastrostomy, and the BMI fell below 18 in eight patients. Regarding the SMA3 patients, one was under NIV for < 16h/day and none had gastrostomy. At T0 and T302, all patients were assessed using standardized motor function tests chosen according to their age and motor function. Functional assessments were performed by expert physiotherapists trained with standardized procedure manuals (66) and reliability sessions. SMA1 patients were assessed with the CHOP-INTEND (67, 68), a functional scale including 16 items aimed at assessing motor function in weak infants. Each item is scored from 0 to 4 (0 being no response and 4 being the complete level of response), with a total score ranging from 0 to 64. SMA2 and SMA3 patients were evaluated with the HFMSE (68, 69), a scale of 33 items investigating the child’s ability to perform different activities. The total score can range from zero, if all the activities are failed, to 66, indicating better motor function. All patients were not wearing spinal jackets or orthoses during the evaluations.

SMA1, SMA2 and SMA3 patients significantly differed in age (SMA1 *vs* SMA2, *P =* 0.006; SMA1 *vs* SMA3, *P* < 0.0001; SMA2 *vs* SMA3, *P =* 0.004; Mann-Whitney test). BMI was lower in SMA1 compared to SMA2 and SMA3 patients (*P =* 0.0001 and *P =* 0.002, respectively; Mann-Whitney test) while sex was not different among SMA groups (χ^2^ = 0.045, *P =* 0.978).

### Intrathecal treatment with Nusinersen

Intrathecal administration of 12 mg of Nusinersen was performed in a hospital environment. Fasting less than 4 h was planned for the procedure in SMA1 patients, while the time between the last meal and the lumbar puncture was 6-8 h in SMA2 and SMA3 patients. In SMA1 the procedure was carried out without sedation, whereas for SMA2 and SMA3 patients a sedation with midazolam was applied. No severe adverse events were reported. After the infusion, all patients were recommended to lie for 2 h to avoid any possible post-lumbar puncture symptoms.

### CSF sample collection

CSF samples were collected at the time of intrathecal administration of Nusinersen in polypropylene tubes and stored at −80°C until further analysis. Amino acid levels were measured in the CSF sample of each patient. Exclusion criteria included the presence of symptoms or changes in blood biochemical and haematological parameters suggestive of a systemic inflammatory state, and/or immunosuppressive treatments ongoing in the last 6 months before inclusion.

### Animals

Experiments in mice were performed according to the international guidelines for animal research and approved by the Animal Care Committee of “Federico II” University of Naples, Italy and the Ministry of Health, Italy. Heterozygous SMNΔ7 carrier mice (Smn^+/-;^ SMN2^+/+^; SMNΔ7^+/+^) were purchased from Jackson Laboratory (stock number 005025) and bred to obtain Smn^+/+^ (WT) animals and Smn^-/-^ (SMA) animals. Mice were housed with 12 h light/dark cycle and were given free access to food and water. All efforts were made to minimize animal suffering and to reduce the number of animals used. The colony was maintained by interbreeding carrier mice, and the offspring were genotyped by PCR assays on tail DNA according to the protocols provided by Jackson Laboratory as previously reported (70). Data were obtained from brain and spinal cord tissue of WT and SMNΔ7 mice isolated at P3 and P11, considering P0 as the day of birth.

### HPLC detection

CSF samples (100 µl) were mixed in a 1:10 dilution with HPLC-grade methanol (900 µl) and centrifuged at 13,000 × g for 10 min; supernatants were dried and then suspended in 0.2 M TCA. Mouse brain and spinal cord frozen samples were homogenized in 1:10 (w/v) 0.2 M TCA, sonicated (4 cycles, 10 s each), and centrifuged at 13,000 × g for 20 min. TCA supernatants from mice and human samples were then neutralized with NaOH and subjected to pre-column derivatization with o-phthaldialdehyde (OPA)/N-acetyl-L-cysteine (NAC). Diasteroisomer derivatives were resolved on a UHPLC Agilent 1290 Infinity (Agilent Technologies, Santa Clara, CA, USA) using a ZORBAX Eclipse Plus C8, 4.6 × 150 mm, 5 μm (Agilent Technologies, Santa Clara, CA, USA) under isocratic conditions (0.1 M sodium acetate buffer, pH 6.2, 1% tetrahydrofuran, and 1.5 mL/min flow rate). A washing step in 0.1 M sodium acetate buffer, 3% tetrahydrofuran, and 47% acetonitrile was performed after every single run. Identification and quantification of D-Asp, L-Asp, L-Glu, D-Ser, L-Ser and L-Gln were based on retention times and peak areas, compared with those associated with external standards. All the precipitated protein pellets from mice samples were solubilized in 1% SDS solution and quantified by bicinchoninic acid (BCA) assay method (Pierce™ BCA Protein Assay Kits, Thermofisher scientific, Rockford, IL, USA). The concentration of amino acids in tissue homogenates was normalized to the total protein content and expressed as nmol/mg protein. Amino acid levels in the CSF were expressed as micromolar (µM).

### Drug treatment and behavioral assays in SMA mice

For studies of D-Ser supplementation in SMA mice, all procedures were performed on postnatal mice in accordance with the NIH guidelines and approved by the Institutional Laboratory Animal Care and Use Committee of Columbia University. FVB.Cg-*Grm7^Tg(SMN2)89Ahmb^ Smn1^tm1Msd^* Tg(SMN2*delta7)4299Ahmb/J (JAX Strain # 005025) mice were interbred to obtain SMA mutant mice (71). Mice were housed in a 12h/12h light/dark cycle with access to food and water *ad libitum*. Mice from all experimental groups were monitored daily for weight, motor function, and survival from birth to 21 days of age. The righting reflex was assessed by placing the mouse on its back and measuring the time it took to turn upright on its four paws (righting time). The cut-off test time was 60s. For each testing session, the test was repeated three times, and the mean of the recorded times was calculated. D-Ser (Sigma #S4250) was dissolved in water, filter sterilized and delivered daily at a dose of 500 mg/kg by intraperitoneal injections starting from P0. Approximately equal proportions of mice of both sexes were used, and aggregated data were presented because gender-specific differences were not found.

### Protein analysis

For Western blot analysis, mice were euthanized and spinal cord collection was performed in a dissection chamber under continuous oxygenation (95%O_2_/5%CO_2_) in the presence of cold (∼12°C) artificial cerebrospinal fluid (aCSF) containing 128.35mM NaCl, 4mM KCl, 0.58mM NaH_2_PO_4_, 21mM NaHCO_3_, 30mM D-Glucose, 1.5mM CaCl_2_, and 1mM MgSO_4_. Total protein extracts were generated by homogenization of spinal cords in SDS sample buffer (2% SDS, 10% glycerol, 5% ß-mercaptoethanol, 60mM Tris-HCl pH 6.8, and bromophenol blue), followed by brief sonication and boiling. Proteins were quantified using the *RC DC*^TM^ Protein Assay (Bio-Rad) and 25µg of protein extract was analyzed by SDS/PAGE on 12% polyacrylamide gels followed by Western blotting as previously described (72). Anti-SMN mouse monoclonal antibody (BD Transd Lab, clone 8, #610646; 1:10,000), anti-GAPDH mouse monoclonal antibody (Sigma, clone 6C5, #MAB374, 1:50,000), and HRP conjugated goat anti-mouse secondary antibody (Jackson #115-035-044; 1:10,000) were used. The signal was detected using an iBrigth CL1500 Imaging System (Thermo Fisher Scientific) and image quantification was processed with the iBright Analysis Software (version 5.1.0).

### Statistical analysis

Statistical analyses were performed using SPSS software v.27 (SPSS Inc., Chicago, IL, USA) and R Language v.4.3.2 (R Foundation for Statistical Computing, Vienna, Austria). Normality distribution was assessed by q-q plot and Shapiro–Wilk test. Quantitative variables were expressed by the median and interquartile range (IQR), while qualitative variables were by absolute or relative frequency. The correlation was evaluated by non-parametric Spearman’s rho. The effect of confounders on correlation was evaluated by partial correlation and multivariate linear regression on natural log-transformed data. Differences between independent groups were studied by the non-parametric Kruskal-Wallis test followed, if statistically significant, by post-hoc tests performed by the Mann-Whitney test with Bonferroni’s correction. The effect of confounders was evaluated by ANCOVA on natural log-transformed variables. Differences between dependent groups were studied by non-parametric Friedman test followed, if statistically significant, by post-hoc tests performed by Wilcoxon Signed Ranks Test with Bonferroni’s correction. Amino acid concentrations in the CNS of SMA and WT mice were compared using Mann-Whitney test and two-way ANOVA followed by Tukey’s *post-hoc* using Prism 8 version 8.0.2. For studies of D-Ser supplementation in SMA mice, statistical analysis of SMN protein levels was performed by one-way ANOVA with Šídák’s multiple comparisons test. Differences in weight gain and motor function were analyzed by two-way ANOVA with Tukey’s multiple comparison test. A comparison of survival curves was performed using the Log-rank (Mantel-Cox) test. Prism 10 for macOS version 10.3.1 was used for these statistical analyses.

## Author Contributions

A.H. and R.d.V. conducted HPLC experiments and acquired data; T.N. acquired data and prepared figures; T.N. and M.V. analyzed data; M.J.C., S.Y. and H.Y. conducted in vivo experiments on SMA mice, acquired and analyzed data; A.D.A., C.P., C.B. and E.B. provided CSF samples of patients; X.K., V.V. and G.P. provided brain and spinal cord samples of SMA mice; F.E., L.P. and A.U. wrote the manuscript; A.U. designed research studies.

## Acknowledgements and Funding

A.U., G.P., T.N., R.d.V., E.B., and A.D.A. were supported by #NEXTGENERATIONEU (NGEU) funded by the Ministry of University and Research (MUR), National Recovery and Resilience Plan (NRRP), project MNESYS (PE0000006) – A Multiscale integrated approach to the study of the nervous system in health and disease (DN. 1553 11.10.2022). E.B. and A.D.A. were also supported by a grant from Ricerca Finalizzata from the Italian Ministry of Health (Project nr RF-2019-12370334); E.B. A.D. and C.B. are members of the ERN NMD European Network (Project nr 2016/557). L.P. was supported by NIH grants R01NS102451, R01NS114218, and R01NS116400.

