## Supplemental material for "Dysregulated balance of D- and L-amino acids modulating glutamatergic neurotransmission in severe spinal muscular atrophy"

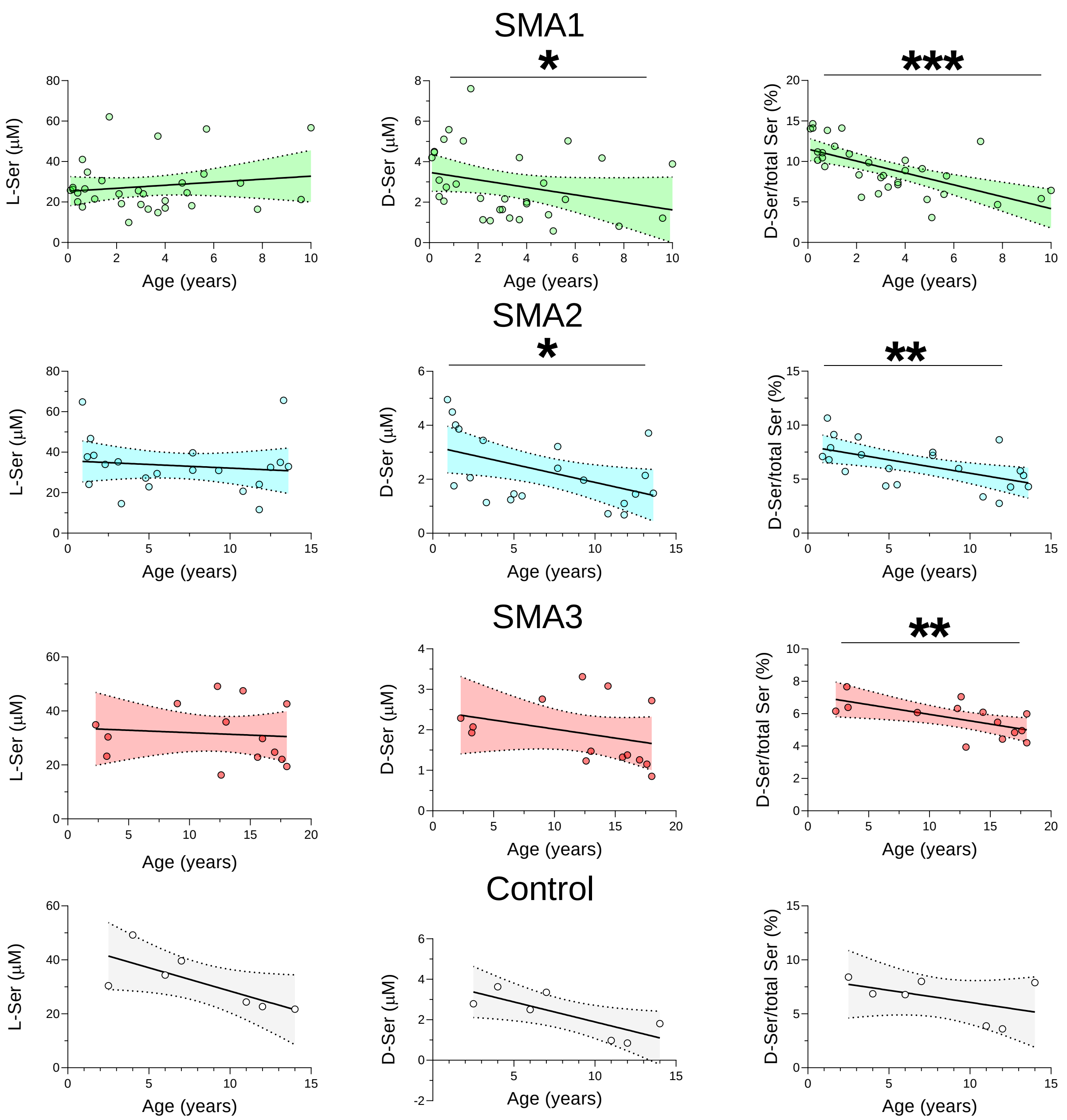


**Suppl Fig 1:** Correlation between age and amino acid concentrations in naive cohorts of SMA1, SMA2 and SMA3 patients as well as normal controls. Association between L-Ser, D-Ser, or D-Ser/tot Ser ratio with age of SMA1, SMA2, SMA3, and control individuals. *$P < 0.05$, **$P < 0.01$, ***$P < 0.0001$, Spearman correlation.

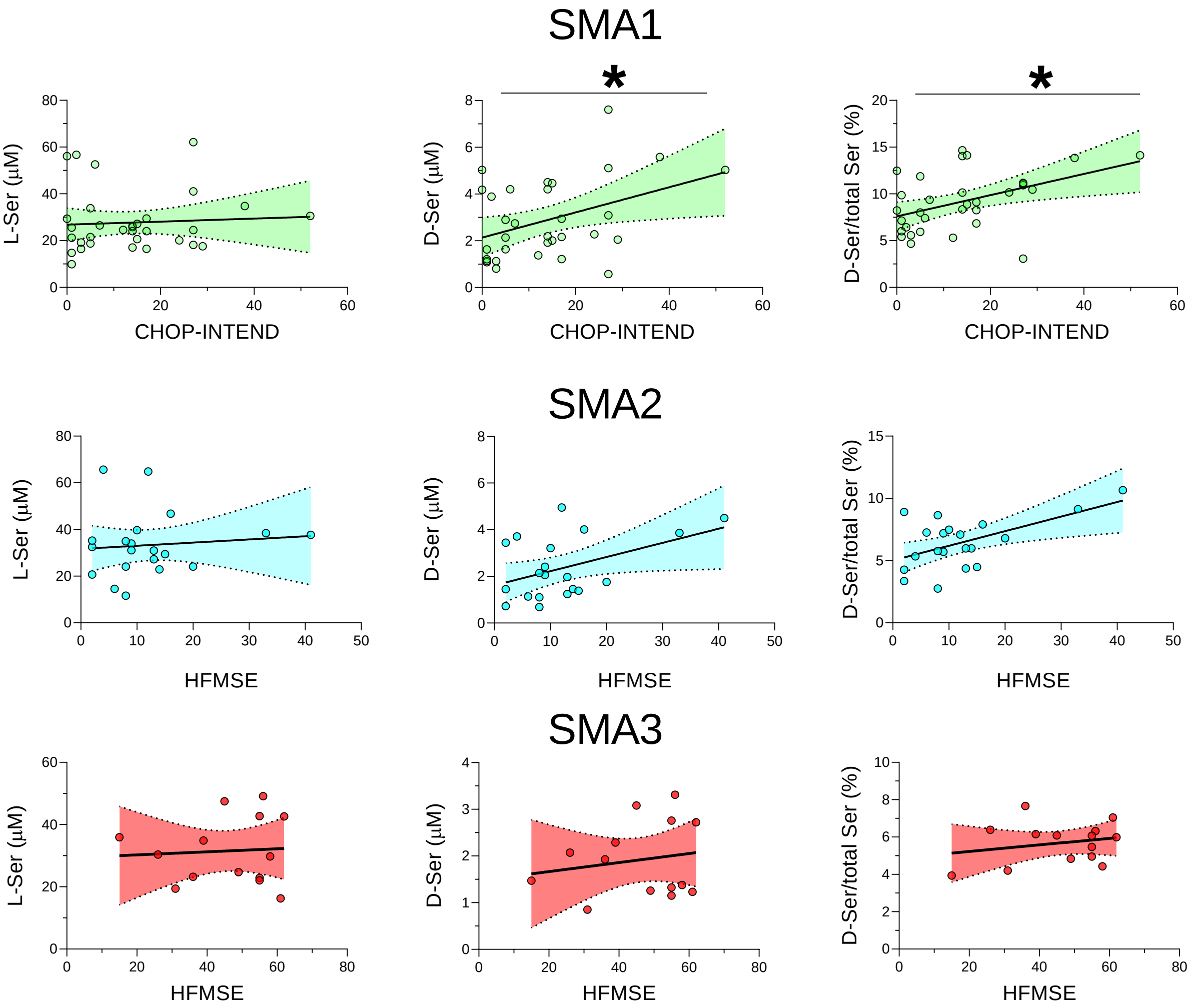


**Suppl Fig 2:** Correlation between amino acid concentrations and CHOP-INTEND or HFMSE in naive cohort of SMA1, SMA2 and SMA3 patients. Association between L-Ser, D-Ser, or D-Ser/tot Ser ratio with CHOP-INTEND (SMA1) or HFMSE (SMA2 and SMA 3). *$P < 0.05$, Spearman correlation.

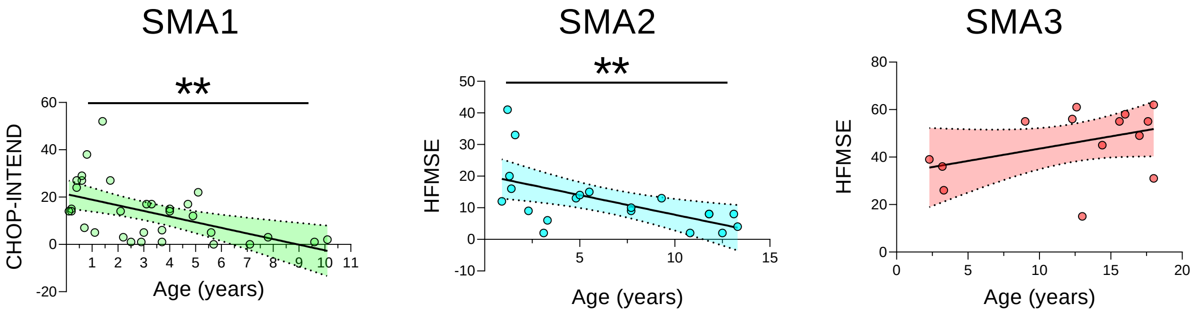


**Suppl Fig 3:** Correlation of CHOP-INTEND or HFMSE with age of naive SMA1, SMA2 and SMA3 patients. $^{**}P < 0.01$, Spearman correlation.

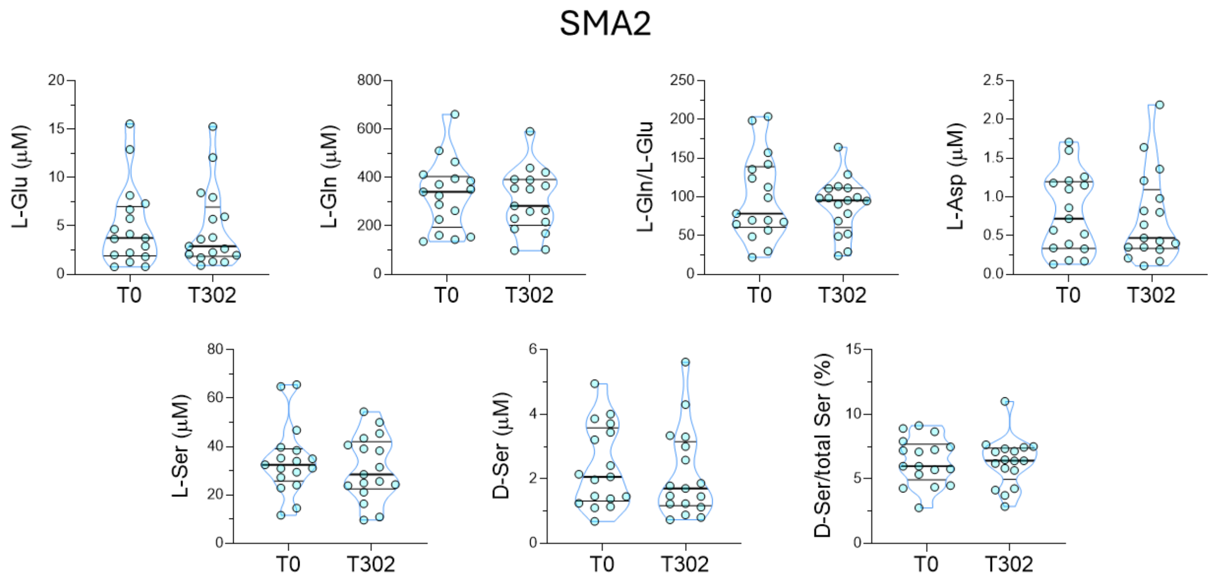


**Suppl Fig 4:** Effect of Nusinersen on amino acids levels in the CSF of SMA2 patients. Levels of L-glutamate (L-Glu), L-glutamine (L-Gln), L-glutamine/L-glutamate ratio (L-Gln/L-Glu), L-aspartate (L-Asp), L-serine (L-Ser), D-serine (D-Ser) and D-serine/total serine (L-Ser/total Ser) percentage in the CSF of SMA2 patients prior to treatment (T0, $n$ = 17) and at the time of the sixth (T302, $n$ = 17) injection of Nusinersen. Data are shown as violin plots representing median with interquartile range (IQR). Dots represent individual patients' values.

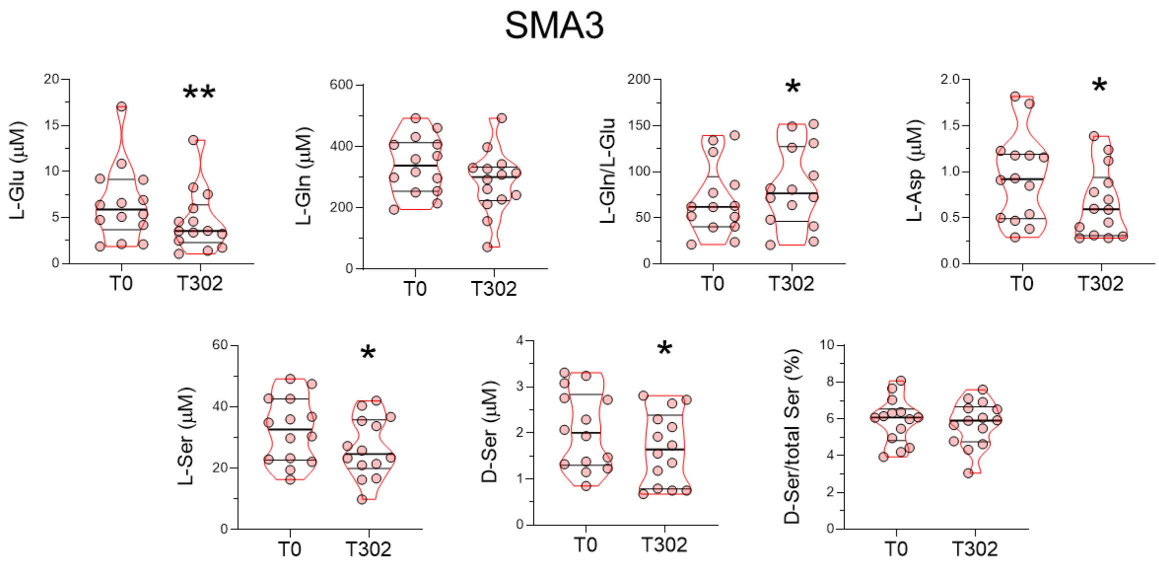


**Suppl Fig 5:** Effect of Nusinersen on amino acids levels in CSF of SMA3 patients. Levels of of L-glutamate (L-Glu), L-glutamine (L-Gln), L-glutamine/L-glutamate (L-Gln/L-Glu) ratio, L-aspartate (L-Asp), L-serine (L-Ser), D-serine (D-Ser) and D-serine/total serine (L-Ser/total Ser) percentage ratio in the CSF of SMA3 patients prior to treatment (T0, $n$ = 14) and at the time of the sixth (T302, $n$ = 14) injection of Nusinersen. *$P < 0.05$, **$P < 0.01$, compared to T0 (Wilcoxon matched-pairs signed ranks test). Data are shown as violin plots representing median with interquartile range (IQR). Dots represent individual patients' values.

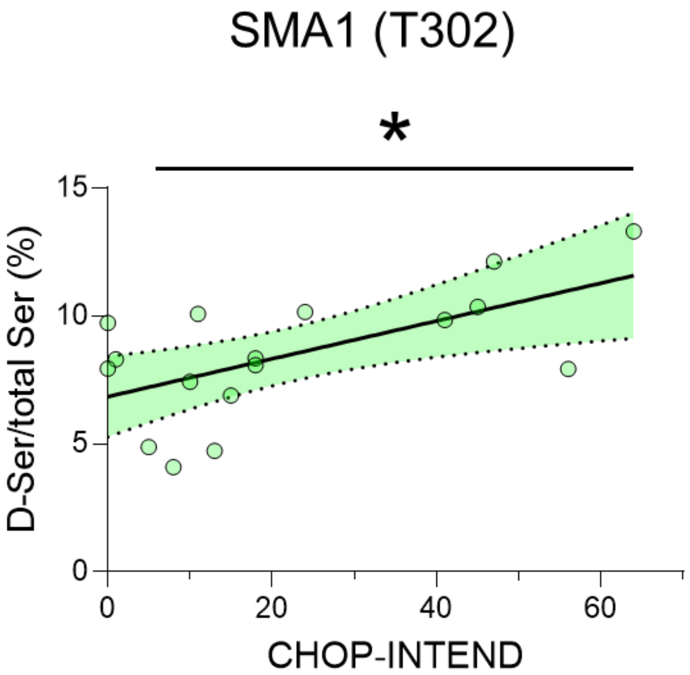


**Suppl Fig 6**: Effect of Nusinersen on the association between D-serine/total serine (L-Ser/total Ser) percentage and CHOP-INTEND of SMA1 patients. *$P < 0.05$, Spearman correlation.

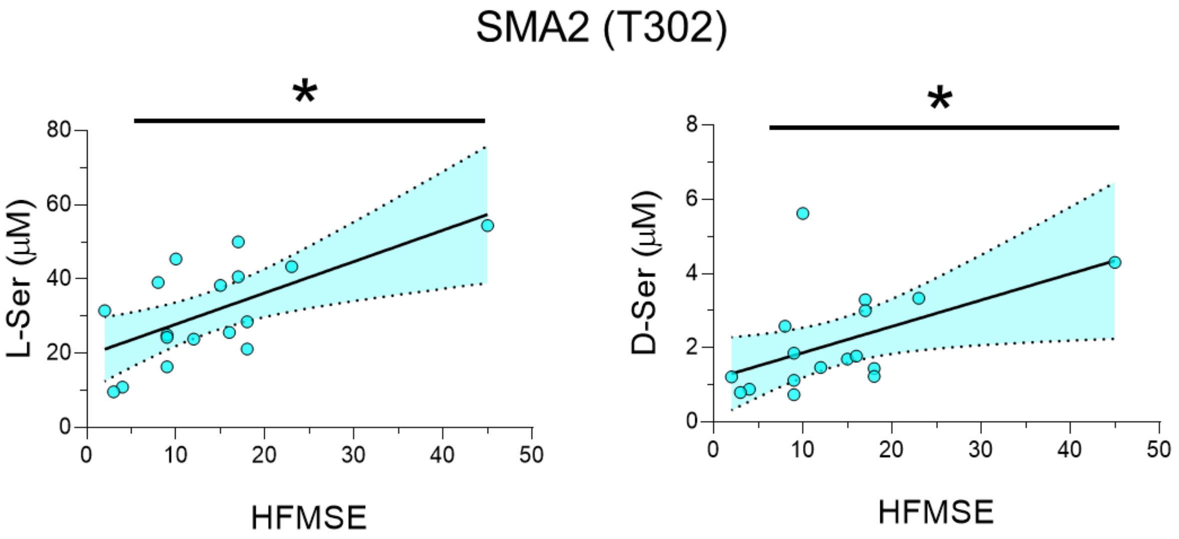


**Suppl Fig 7:** Effect of Nusinersen on the association between L-Ser or D-Ser and HFMSE of SMA2 patients. *$P < 0.05$, Spearman correlation.

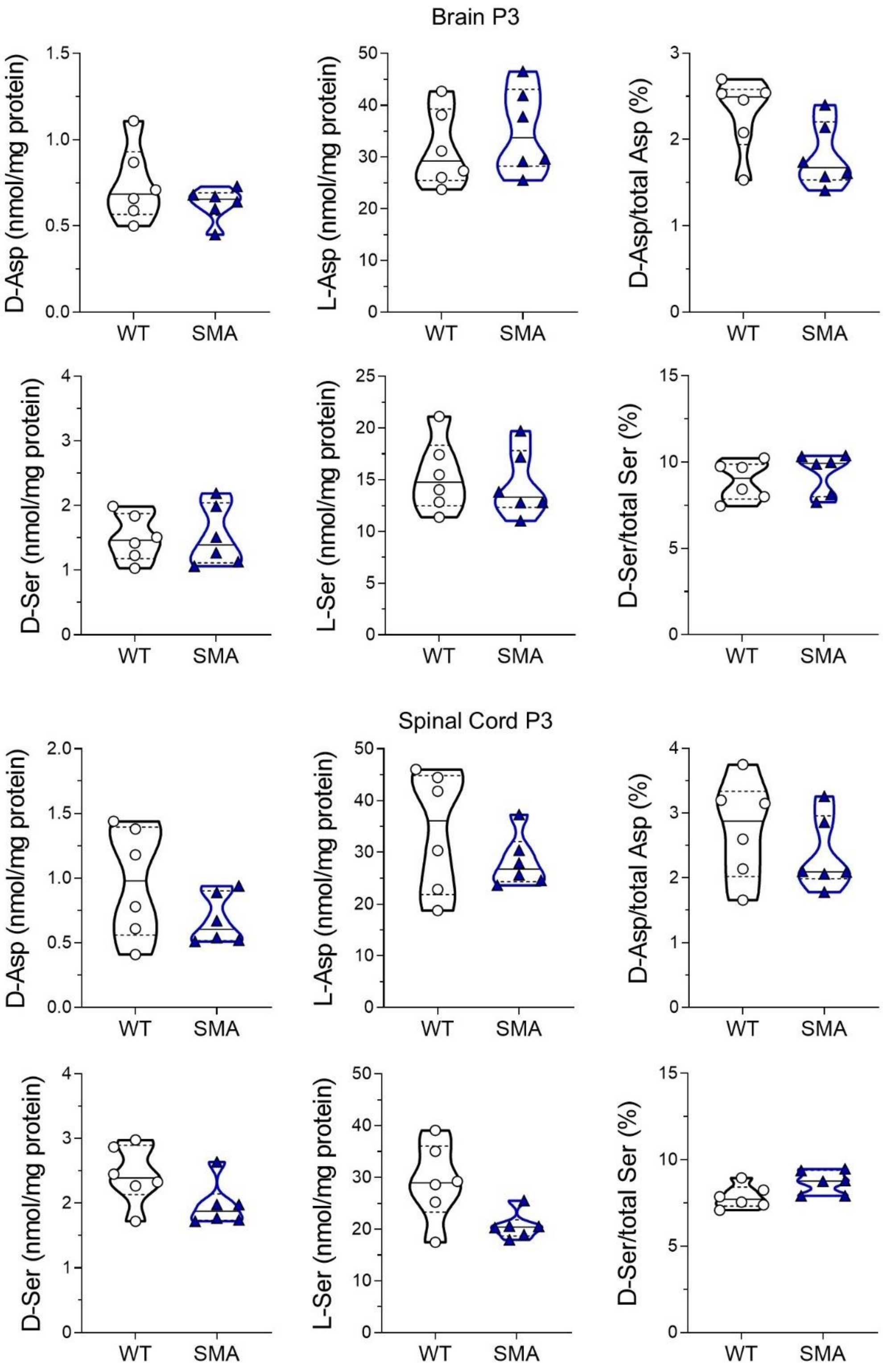


**Suppl Fig 8:** Analysis of D-aspartate (D-Asp), L-aspartate (L-Asp), D-serine (D-Ser) and L-serine (L-Ser) and their relative ratios between WT and SMA mice in the brain and spinal cord at P3. The average amounts of amino acids detected were normalized for mg of proteins. Dots represent values from individual mice. Amino acids levels are expressed as violin plots representing median with interquartile range (IQR) and analyzed by Mann-Whitney.

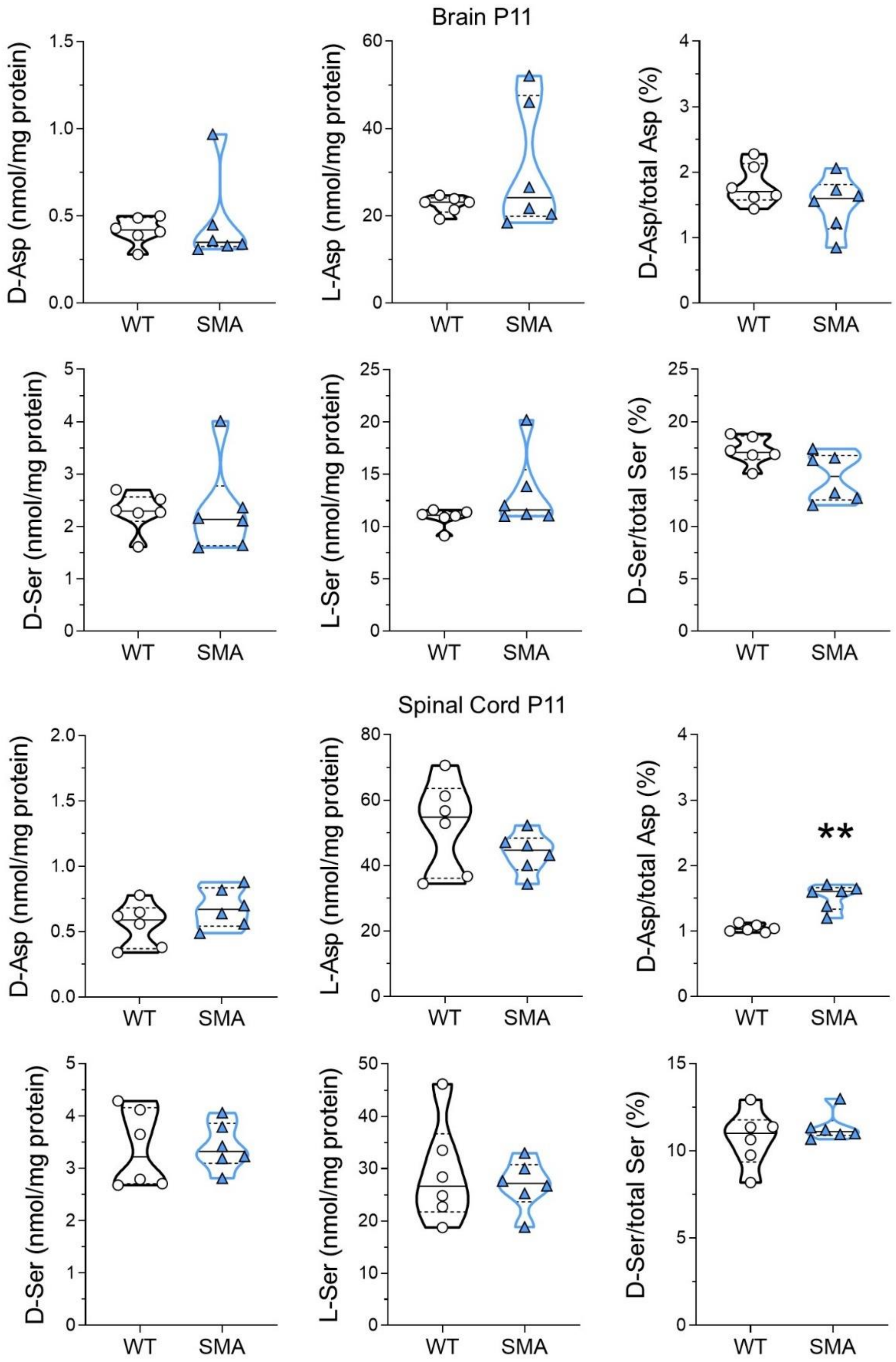


**Suppl Fig 9:** Analysis of D-aspartate (D-Asp), L-aspartate (L-Asp), D-serine (D-Ser) and L-serine (L-Ser) and their relative ratios between WT and SMA mice in the brain and spinal cord at P11. The average amounts of amino acids detected were normalized for mg of proteins. Dots represent values from individual mice. Amino acids levels are expressed as violin plots representing median with interquartile range (IQR) and analyzed by Mann-Whitney. $^{**}P < 0.01$, compared to age-matched WT mice.

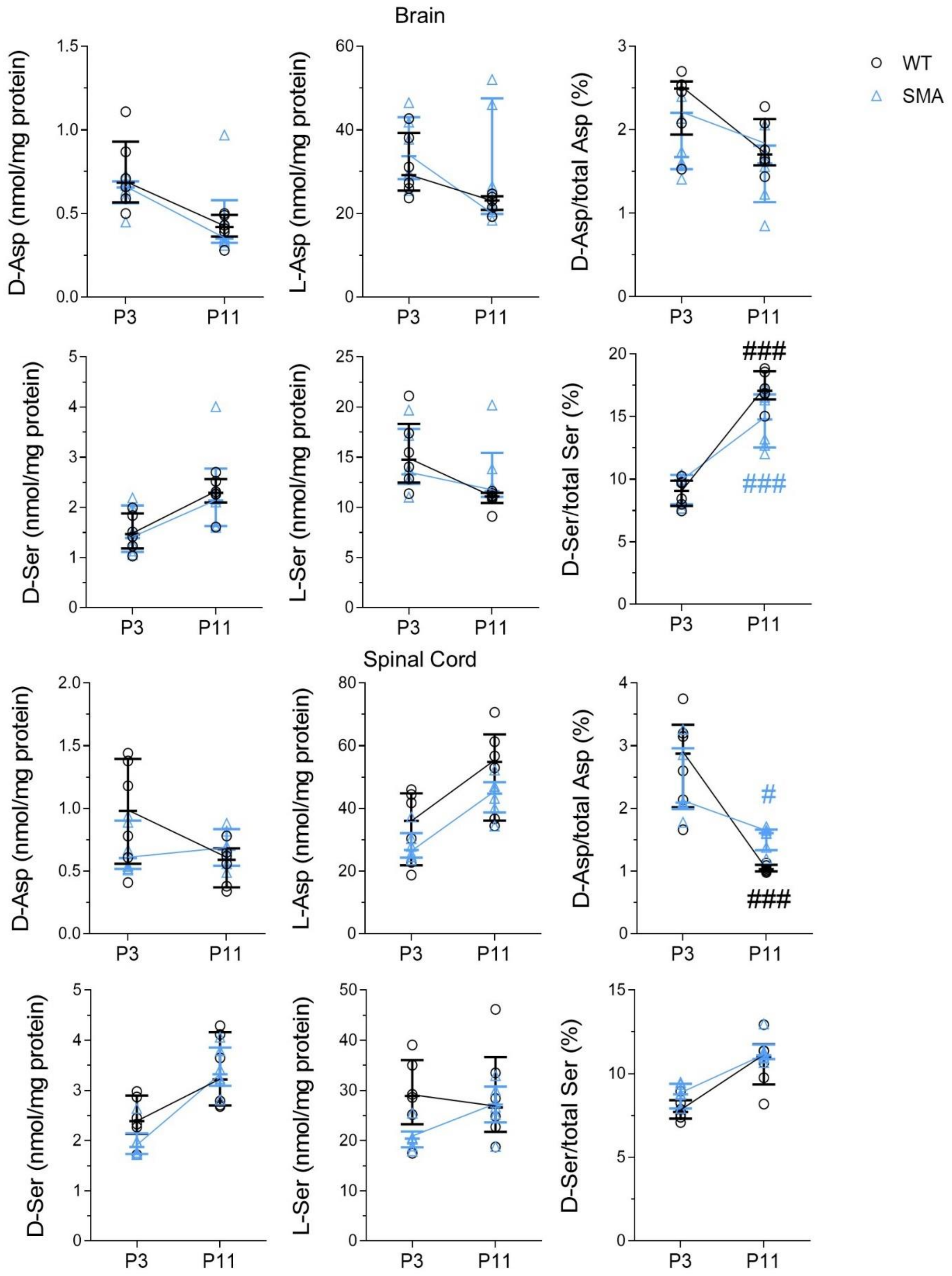


**Suppl Fig 10:** Analysis of D-aspartate (D-Asp), L-aspartate (L-Asp), D-serine (D-Ser) and L-serine (L-Ser) and their relative ratios between WT and SMA mice at P3 and P11 analyzed by two-way ANOVA followed by Tukey's multiple comparisons test. #*P* < 0.05, ##*P* < 0.01, P11 compared to genotype-matched P3 mice. Amino acids levels were shown as scatter dot plots representing median with interquartile range (IQR) while dots represent individual animals' values.

**Supplementary Table 1.** Amino acids levels in controls and SMA patients.

| Amino acids | Controls (*n* = 7) | SMA1 (*n* = 34) | SMA2 (*n* = 22) | SMA3 (*n* = 17) | Kruskal-Wallis (*P*-value)[a] | Post-hoc analysis (*P*-value)[b] |
|---|---|---|---|---|---|---|
| L-Gln (μM) | 257 (206-315) | 256 (174-288) | 307 (211-400) | 312 (259-406) | **0.023** | SMA1 vs SMA3: **0.024** |
| L-Glu (μM) | 8.96 (5.28-9.34) | 2.50 (1.11-5.04) | 3.96 (2.15-6.86) | 5.36 (2.29-9.16) | **0.007** | Controls vs SMA1: **0.024** |
| L-Gln/L-Glu | 27.5 (21.7-59.7) | 120 (53.4-188) | 74.2 (54.9-129) | 62.3 (40.1-119) | **0.005** | Controls vs SMA1: **0.006**<br>Controls vs SMA2: **0.036** |
| L-Asp (μM) | 0.83 (0.40-0.87) | 0.39 (0.23-0.67) | 0.63 (0.34-1.16) | 0.85 (0.43-1.18) | **0.037** | |
| L-Ser (μM) | 30.4 (22.7-39.6) | 25.0 (19.1-34.1) | 32.7 (24.1-38.7) | 29.8 (23.1-39.7) | 0.166 | |
| D-Ser (μM) | 2.50 (0.98-3.36) | 2.47 (1.57-4.20) | 2.01 (1.35-3.54) | 1.93 (1.26-2.74) | 0.374 | |
| D-Ser/total Ser (%) | 6.86 (3.88-8.01) | 8.61 (6.74-11.1) | 6.39 (4.45-7.73) | 6.07 (4.63-6.71) | <**0.001** | SMA1 vs SMA2: **0.012**<br>SMA1 vs SMA3: <**0.001** |

Data are expressed as median (IQR). Analysis by non-parametric Kruskal-Wallis[a], post-hoc analysis by Mann-Whitney with Bonferroni's correction[b]. L-Gln, L-Glu, L-Gln/L-Glu ratio, L-Asp, D-Ser/total Ser (%) levels were natural log transformed and ANCOVA analysis was performed, considering sex and age as possible confounders. At ANCOVA, only L-Glu ($P$ = 0.006) and L-Glu/L-Gln ratio ($P$ = 0.003) remain significantly associated. Abbreviation: L-glutamine (L-Gln), L-glutamate (L-Glu), L-glutamine/L-glutamate (L-Gln/L-Glu), L-aspartate (L-Asp), L-serine (L-Ser), D-serine (D-Ser), D-serine/total serine (D-Ser/total Ser).

**Supplementary Table 2.** Variations of amino acids at T0 vs T302 among SMA patients.

| Amino acids | SMA1<br>T0 vs T302 | SMA2<br>T0 vs T302 | SMA3<br>T0 vs T302 |
|---|---|---|---|
| L-Gln (μM) | 237 vs 285 (***P* = 0.025**) (*n* = 18) | 341 vs 283 (*P* = 0.758) (*n* = 17) | 338 vs 300 (*P* = 0.064) (*n* = 14) |
| L-Glu (μM) | 2.80 vs 3.70 (***P* = 0.035**) (*n* = 18) | 3.76 vs 2.91 (*P* = 0.831) (*n* = 17) | 5.87 vs 3.54 (***P* = 0.006**) (*n* = 14) |
| L-Gln/L-Glu | 92.1 vs 102 (*P* = 0.777) (*n* = 18) | 78.4 vs 95.5 (*P* = 0.492) (*n* = 17) | 62.0 vs 76.7 (***P* = 0.019**) (*n* = 14) |
| L-Asp (μM) | 0.37 vs 0.57 (*P* = 0.287) (*n* = 18) | 0.72 vs 0.47 (*P* = 0.379) (*n* = 17) | 0.92 vs 0.59 (***P* = 0.038**) (*n* = 14) |
| L-Ser (μM) | 24.5 vs 35.4 (***P* = 0.004**) (*n* = 18) | 32.5 vs 28.5 (*P* = 0.586) (*n* = 17) | 32.6 vs 24.6 (***P* = 0.019**) (*n* = 14) |
| D-Ser (μM) | 2.82 vs 2.83 (*P* = 0.248) (*n* = 18) | 2.06 vs 1.70 (*P* = 0.492) (*n* = 17) | 2.00 vs 1.64 (***P* = 0.016**) (*n* = 14) |
| D-Ser/total Ser (%) | 9.24 vs 8.19 (***P* = 0.007**) (*n* = 18) | 5.98 vs 6.41 (*P* = 0.653) (*n* = 17) | 6.08 vs 5.91 (*P* = 0.221) (*n* = 14) |

Data are expressed as median. *P*-values are related to non-parametric Wilcoxon test. The number (*n*) of patients is also indicated.

**Supplementary Table 3.** Levels of amino acids in the brain and spinal cord of wild type (WT) and SMNΔ7 mice at P3 and P11.

| | | BRAIN | | | | SPINAL CORD | | | |
|---|---|---|---|---|---|---|---|---|---|
| | | P3 | | P11 | | P3 | | P11 | |
| | | WT | SMA | WT | SMA | WT | SMA | WT | SMA |
| L-Glu | Median | 69.33 | 71.51 | 71.92 | 67.07 | 107.0 | 77.42 | 133.5 | 107.5 |
| | (IQR) | (59.58-88.42) | (54.71-86.47) | (64.26-73.90) | (53.54-85.03) | (78.22-129.5) | (72.51-87.12) | (103.7-151.8) | (94.23-125.8) |
| | *P*-value | 0.5887 | | 0.5887 | | 0.0931 | | 0.1797 | |
| L-Gln | Median | 48.58 | 48.61 | 37.21 | 50.09 | 274.3 | 209.8 | 152.6 | 220.3 |
| | (IQR) | (45.17-70.35) | (40.17-54.23) | (32.06-41.73) | (46.70-62.42) | (205.4-324.2) | (194.7-216.0) | (126.1-185.6) | (192.5-255.8) |
| | *P*-value | 0.5887 | | **0.0043** | | 0.0649 | | **0.0152** | |
| L-Gln/L-Glu | Median | 0.750 | 0.620 | 0.560 | 0.740 | 2.590 | 2.585 | 1.230 | 1.995 |
| | (IQR) | (0.663-0.875) | (0.608-0.753) | (0.468-0.633) | (0.728-0.893) | (2.500-2.678) | (2.500-2.750) | (1.070-1.313) | (1.763-2.295) |
| | *P*-value | 0.2900 | | **0.0022** | | 0.9697 | | **0.0022** | |
| D-Ser | Median | 1.465 | 1.390 | 2.290 | 2.135 | 2.390 | 1.875 | 3.220 | 3.325 |
| | (IQR) | (1.180-1.878) | (1.113-2.040) | (2.098-2.565) | (1.630-2.773) | (2.133-2.898) | (1.735-2.145) | (2.703-4.163) | (3.095-3.858) |
| | *P*-value | 0.9091 | | 0.4848 | | 0.1429 | | 0.8182 | |
| L-Ser | Median | 14.76 | 13.31 | 11.10 | 11.61 | 28.93 | 20.40 | 26.63 | 27.18 |
| | (IQR) | (12.48-18.34) | (12.33-17.82) | (10.44-11.45) | (11.04-15.44) | (23.28-36.04) | (18.67-21.81) | (21.73-36.67) | (23.65-90.76) |
| | *P*-value | 0.4848 | | 0.1797 | | 0.0931 | | >0.9999 | |
| D-Ser/total Ser | Median | 9.065 | 9.935 | 17.07 | 14.78 | 7.715 | 8.770 | 11.01 | 11.11 |
| | (IQR) | (7.865-9.880) | (8.003-10.34) | (16.38-18.64) | (12.54-16.77) | (7.323-7.920) | (8.425-9.398) | (9.37-11.77) | (10.88-11.75) |
| | *P*-value | 0.3095 | | 0.0649 | | 0.0584 | | 0.6991 | |
| D-Asp | Median | 0.685 | 0.655 | 0.420 | 0.350 | 0.980 | 0.605 | 0.590 | 0.670 |
| | (IQR) | (0.567-0.930) | (0.563-0.693) | (0.363-0.493) | (0.325-0.580) | (0.560-1.395) | (0.518-0.903) | (0.370-0.683) | (0.543-0.835) |
| | *P*-value | 0.5887 | | 0.5887 | | 0.3095 | | 0.2576 | |
| L-Asp | Median | 29.25 | 33.73 | 23.13 | 24.16 | 36.08 | 26.76 | 54.85 | 44.71 |
| | (IQR) | (25.47-39.28) | (28.24-43.07) | (20.89-24.15) | (19.94-47.58) | (21.83-44.83) | (24.33-32.09) | (36.15-63.62) | (38.68-44.71) |
| | *P*-value | 0.5887 | | 0.6667 | | 0.5887 | | 0.2403 | |
| D-Asp/total Asp | Median | 2.495 | 1.675 | 1.705 | 1.600 | 2.875 | 2.095 | 1.030 | 1.605 |
| | (IQR) | (1.943-2.580) | (1.530-2.205) | (1.575-2.130) | (1.135-1.813) | (2.020-3.338) | (1.990-2.960) | (0.995-1.100) | (1.335-1.665) |
| | *P*-value | 0.0931 | | 0.2403 | | 0.3939 | | **0.0022** | |

Data are expressed as median. *P*-values are related to non-parametric Mann-Whitney test, $n = 6$ animals per group.

**Supplementary Table 4.** Age effect on amino acids levels in the brain and spinal cord of wild type (WT) and SMA mice at P3 and P11.

| | | BRAIN | | | SPINAL CORD | | |
|---|---|---|---|---|---|---|---|
| | | **F (DFn, DFd)** | ***P*-value** | **Tukey's multiple comparisons test** | **F (DFn, DFd)** | ***P*-value** | **Tukey's multiple comparisons test** |
| L-Glu | Interaction | F (1, 20) = 0.1622 | 0.6914 | | F (1, 20) = 0.029 | 0.8657 | |
| | Age | F (1, 20) = 0.1019 | 0.7529 | | F (1, 20) = 10.82 | 0.0037 | |
| | Genotype | F (1, 20) = 0.02570 | 0.8742 | | F (1, 20) = 6.201 | 0.0217 | |
| L-Gln | Interaction | F (1, 20) = 6.632 | **0.0181** | P3:SMA vs. P3:WT *P* = 0.6519<br>P3:SMA vs. P11:SMA *P* = 0.6444<br>P3:WT vs. P11:WT *P* = 0.0978<br>P11:SMA vs. P11:WT *P* = 0.0955 | F (1, 20) = 13.45 | **0.0015** | P3:SMA vs. P3:WT *P* = 0.0970<br>P3:SMA vs. P11:SMA *P* = 0.8975<br>P3:WT vs. P11:WT *P* = **0.0012**<br>P11:SMA vs. P11:WT *P* = 0.0583 |
| | Age | F (1, 20) = 0.8153 | 0.3773 | | F (1, 20) = 7.202 | 0.0143 | |
| | Genotype | F (1, 20) = 0.8474 | 0.3682 | | F (1, 20) = 0.03362 | 0.8564 | |
| L-Gln/L-Glu | Interaction | F (1, 20) = 14.71 | **0.001** | P3:SMA vs. P3:WT *P* = 0.4531<br>P3:SMA vs. P11:SMA *P* = 0.2055<br>P3:WT vs. P11:WT *P* = **0.0145**<br>P11:SMA vs. P11:WT *P* = **0.0043** | F (1, 20) = 20.21 | **0.0002** | P3:SMA vs. P3:WT *P* > 0.9999<br>P3:SMA vs. P11:SMA *P* = **0.0028**<br>P3:WT vs. P11:WT *P* < **0.0001**<br>P11:SMA vs. P11:WT *P* < **0.0001** |
| | Age | F (1, 20) = 0.8907 | 0.3566 | | F (1, 20) = 106.2 | <0.0001 | |
| | Genotype | F (1, 20) = 2.913 | 0.1033 | | F (1, 20) = 19.89 | 0.0002 | |
| D-Ser | Interaction | F (1, 20) = 8.407E-4 | 0.9772 | | F (1, 20) = 1.440 | 0.2442 | |
| | Age | F (1, 20) = 11.56 | 0.0028 | | F (1, 20) = 31.61 | <0.0001 | |
| | Genotype | F (1, 20) = 0.01518 | 0.9032 | | F (1, 20) = 0.9909 | 0.3314 | |
| L-Ser | Interaction | F (1, 20) = 1.670 | 0.211 | | F (1, 20) = 1.317 | 0.2647 | |
| | Age | F (1, 20) = 5.610 | 0.028 | | F (1, 20) = 1.284 | 0.2706 | |
| | Genotype | F (1, 20) = 0.3861 | 0.5414 | | F (1, 20) = 3.733 | 0.0676 | |
| D-Ser/total Ser | Interaction | F (1, 20) = 5.412 | **0.0306** | P3:SMA vs. P3:WT *P* = 0.9557<br>P3:SMA vs. P11:SMA *P* < **0.0001**<br>P3:WT vs. P11:WT *P* **< 0.0001**<br>P11:SMA vs. P11:WT *P* = 0.052 | F (1, 20) = 0.06080 | 0.8077 | |
| | Age | F (1, 20) = 112.4 | <0.0001 | | F (1, 20) = 43.27 | <0.0001 | |
| | Genotype | F (1, 20) = 2.576 | 0.1242 | | F (1, 20) = 3.189 | 0.0893 | |
| D-Asp | Interaction | F (1, 20) = 1.118 | 0.3029 | | F (1, 20) = 3.824 | 0.0646 | |
| | Age | F (1, 20) = 11.25 | 0.0032 | | F (1, 20) = 3,702 | 0.0687 | |
| | Genotype | F (1, 20) = 0.2173 | 0.6461 | | F (1, 20) = 0,580 | 0.4551 | |
| L-Asp | Interaction | F (1, 20) = 0.3941 | 0.5372 | | F (1, 20) = 0.089 | 0.7685 | |
| | Age | F (1, 20) = 3.065 | 0.0953 | | F (1, 20) = 17.09 | 0.0005 | |
| | Genotype | F (1, 20) = 2.503 | 0.1293 | | F (1, 20) = 2.962 | 0.1007 | |
| D-Asp/total Asp | Interaction | F (1, 20) = 0.4019 | 0.5333 | | F (1, 20) = 4.789 | **0.0407** | P3:SMA vs. P3:WT *P* = 0.5209<br>P3:SMA vs. P11:SMA *P* = **0.0363**<br>P3:WT vs. P11:WT *P* < **0.0001**<br>P11:SMA vs. P11:WT *P* = 0.3463 |
| | Age | F (1, 20) = 6.351 | 0.0203 | | F (1, 20) = 40.51 | <0.0001 | |
| | Genotype | F (1, 20) = 6.142 | 0.0222 | | F (1, 20) = 0.05086 | 0.8239 | |

Data were analyzed by two-way ANOVA, followed by Tukey's multiple comparisons test.
